# Compensating functional connectivity changes due to structural connectivity damage via modifications of local dynamics

**DOI:** 10.1101/2024.05.31.596792

**Authors:** Sophie Benitez Stulz, Samy Castro, Gregory Dumont, Boris Gutkin, Demian Battaglia

## Abstract

Neurological pathologies as e.g. Alzheimer’s Disease or Multiple Sclerosis are often associated to neurodegenerative processes affecting the strength and the transmission speed of long-range inter-regional fiber tracts. Such degradation of Structural Connectivity impacts on large-scale brain dynamics and the associated Functional Connectivity, eventually perturbing network computations and cognitive performance. Functional Connectivity however is not bound to merely mirror Structural Connectivity, but rather reflects the complex coordinated dynamics of many regions. Here, using analytical characterizations of toy models and computational simulations connectome-base whole-brain models, we predict that suitable modulations of regional dynamics could precisely compensate for the effects of structural degradation, as if the original Structural Connectivity strengths and speeds of conduction were effectively restored. The required dynamical changes are widespread and aspecific (i.e. they do not need to be restricted to specific regions) so that they could be potentially implemented via neuromodulation or pharmacological therapy, globally shifting regional excitability and/or excitation/inhibition balance. Computational modelling and theory thus suggest that, in the future therapeutic interventions could be designed to “repair brain dynamics” rather than structure to boost functional connectivity without having to block or revert neurodegenerative processes.

**AUTHOR SUMMARY:** Neurological disorders affect Structural Connectivity, i.e. the wiring infrastructure interlinking distributed brain regions. Here we propose that the resulting disruptions in Functional Connectivity, i.e. inter-regional coordination and information sharing, could be compensated by modifying local dynamics so to effectively emulate the restoration of Structural Connectivity (but through a suitable “software patch” rather than by repairing the “hardware”). For simple toy models involving a few regions we can achieve an analytical understanding of how structural and dynamical changes jointly control Functional Connectivity. We then show that the concept of “effective connectome change” via modulation of dynamics robustly extend also to simulation of large-scale models embedding realistic whole-brain connectivity. We thus forecast that novel therapeutic strategies could be devised, targeting dynamics rather than neurodegenerative mechanisms.

## INTRODUCTION

Structural Connectivity (SC) between different brain describes anatomical connections between neuronal populations and regions and, when systematically reconstructed for specific brain subsystems or even complete brains, can be compiled into matrices of large-scale connectivity known as structural connectomes (Sporns et al., 2005; Hagmann et al., 2008; Markov et al., 2014). On the other hand, Functional Connectivity (FC) is meant to describe interactions and information exchange or integration between distinct regions and is estimated through suitable data-driven metrics computed from multivariate time-series of neural activity (Park & Friston, 2013). For instance, zero-lag correlation between dynamic fluctuations is often used as a plain measure of FC (Fox & Raichle, 2007), especially in the case of neural activity with a strong oscillatory component, for which close phase synchronization provides an actual mechanism facilitating information transmission (Fries, 2005; Bastos et al., 2014). Although defined in terms of independent notions, it is natural to suppose that SC strongly contributes to shape the observed FC. Indeed, SC and FC estimated from resting state fluctuations of BOLD activity are highly correlated (Honey et al., 2007; 2009; Greicius et al., 2009). Weakened or lesioned inter-regional SC leads in many cases to weakened FC as well (Alstott et al., 2009). Yet, FC is not identical to SC. It is usually associated to denser graphs, indicating the possibility of coordination even between poorly connected sites. Furthermore, FC can reorganize flexibly following SC damage (Achard et al., 2012), indicating the existence of compensation mechanisms and potential FC homeostasis. Here, using a combination of theory and computational simulations, we will demonstrate the possibility to modulate local dynamics within regions to preserve FC despite damages of increasing severity to the underlying SC infrastructure.

Eventually, theoretical works have suggested that “function follows dynamics” (rather than “structure”). FC is not a mere copy of SC but rather a manifestation of the dynamical states of the system, constrained, but not fully determined by the underlying SC (Kirst et al., 2016). To a multiplicity of dynamical states existing alternatively on top of a same SC connectome can thus correspond a multiplicity of possible FC states (Battaglia et al., 2012; Dumont & Gutkin, 2019), a phenomenon that has been called “functional multiplicity” (from one SC to many FCs; Battaglia & Brovelli, 2020). Conversely, different SCs able to support nevertheless equivalent dynamical states could give rise to essentially equivalent FCs (Stetter et al., 2012; Kispersky et al., 2011), a phenomenon denoted as “structural degeneracy” (from many SCs to a common FC; Battaglia & Brovelli, 2020). This means that changes observed at the level of the functional connectome may be due to changes occurring at the level of the “dynome” –i.e. the set of possible dynamical states of a system (Kopell et al., 2014)– and not only at the level of the connectome. This indirect nature of the link between SC and FC makes its investigation more difficult and potentially requesting the modelling of SC-constrained dynamics to predict FC (Arbabyazd et al., 2021). However, it also enables possible mechanisms –of dynamical, rather than structural nature– for the control of FC on top of an unchanged SC.

Specifically, aging or neurodegenerative diseases, such as e.g. AD or ALS, led to widespread degenerations of long-range SC, with a loss in connection fibers (Salat, 2011; Bateman et al., 2012; Delbeuck et al, 2013; reduced strength of SC links) or a demyelination of axons (Peterson & Fujinami, 2007; Papuć & Rejdak, 2018; You et al., 2019; reduced speed of conduction of SC links). Such damages of SC are paralleled by substantial alterations of FC (Supekar et al., 2008; Dennis & Thompson, 2014). However, the functional deficits induced by a fixed amount of SC degeneration may strongly differ across subjects and patients. The fact that some subjects may maintain normal performance levels despite SC degeneration is often explained in terms of the existence of a so-called “cognitive reserve” (Snowdon et al., 2003; Rentz et al., 2010) and activity-level compensative mechanisms (Bondi et al., 2005). However, the neural basis of such compensation is still poorly understood. Here we hypothesize that maintenance of a “healthy-like” dynamical state allows the preservation of FC organization, despite SC degeneration. Variations of SC may push the system out of the correct dynamic working point of operation. Nevertheless, global, unspecific variations throughout the system of regional features, such as excitatory-inhibitory balance or inhibitory strength, may modify local dynamics and do so in a way to compensate for opposite variations induced by SC changes. In this way the system would be maintained (or re-enter) into the right dynamic regime and the associated FC re-instated. We would thus assist to a net *compensation by dynamics* of FC changes induced by SC degeneration, as if the SC had been effectively “repaired” (which is not the case).

To explore the plausibility of such a scenario, we resort here to computational modelling of systems composed of many coupled regions, each exhibiting oscillatory dynamics. As previously mentioned, flexible patterns of oscillatory synchronization have been proposed as a device to achieve a flexible FC on top of a static SC, by the control of inter-regional phase relations. According to the “Communication-Through-Coherence” hypothesis, two regions with a phase-lag close to in-phase can exchange more easily information than regions whose oscillations are separated by larger phase differences up to anti-phase (Bastos et al., 2014). Such hypothesis is supported both by empirical evidence (Gregoriou et al., 2009; Grothe et al., 2018) and computational analyses (Battaglia et al., 2012; Palmigiano et al., 2017). Different mechanisms for the local generation of oscillations have been proposed, such as the “Pyramidal-Inhibitory-Network-Generated” (PING) or the “Inhibitory-Network-Generated” (ING) scenarios (Whittington et al., 2000; Tiesinga & Sejnowsky, 2009). Their reproduction in mathematical models allows understanding how neuronal-level properties, such as the strengths of excitatory and inhibitory couplings within a region, affect the properties of the collective oscillation of the local region. By using neural mass descriptions of these collective oscillations, it is possible to analytically predict which functional connectivity motifs –corresponding to stable phase-locked configurations (Battaglia et al., 2012; Palmigiano et al., 2017; Dumont & Gutkin, 2019)– can exist among weakly coupled oscillating populations. An important theoretical prediction is thus that the stable phase difference between coupled neuronal populations should not depend exclusively on the properties of the long-range coupling between regions, but also on the properties of local dynamics within regions and, notably, on the macroscopic *phase response curve* (PRC) of their collective oscillations (Dumont et al., 2017; 2019).

Here we will show that modifications of local properties, such as balance between excitatory and inhibitory couplings or the strength of recurrent inhibition, can produce PRC alterations precisely compensating for the alteration of long-range SC, so that the same phase relations as prior to SC degeneration can be re-established. Firstly, we will consider simple toy models of two interconnected regions, for which a full analytical understanding of the SC-to-FC mapping via dynamics can be achieved, for both the PING and ING scenarios for local oscillatory generation. Secondly, we will confirm via numerical simulations that the results obtained in these simple models generalize to whole-brain connectome-based models, involving tens of regions, coupled by anatomically realistic SC connectome matrices.

Our models, although not yet adapted to describe specific neurodegenerative pathologies, already provide a proof-of-concept of the possibility of effective compensation by dynamics of SC-damage-induced alterations of FC. We hypothesize that this general phenomenon could contribute to natural mechanisms for the resilient preservation of FC, eventually mediated by neuromodulatory changes (Backman et al., 2006). In perspective, we also speculate that it could be activated by ad hoc, model-optimized pharmacological interventions aiming at modulating local dynamics to restore global FC (see *Discussion*).

## RESULTS

### Models of PING and ING oscillations

As mentioned in the Introduction, we focus on modelling FC between oscillating regions. The first step is thus constructing models of the local generation of oscillations between each region. Two main scenarios have been described in the literature to explain the emergence of oscillatory activity from the interplay of excitatory (E) and inhibitory (I) populations. In a first “Pyramidal-Inhibitory-Network-Generated” (PING) scenario, growing excitatory (E) activity drives enhanced activation of inhibitory (I) neurons which on their turn reduces excitatory activity, followed by reduced inhibition, in a cyclic manner. In a second “Inhibitory-Network-Generated” (ING) scenario, the strong increase of inhibitory activity prevents any neuron in the network to fire until when inhibitory activity itself is silenced, removing this inhibitory blockade. However, since inhibitory action is not instantaneous, there are tight windows of time, periodically spaced, in which both excitatory and inhibitory neurons can fire. This case, in which only inhibitory interneurons are responsible for the generation of the rhythm and the excitatory neurons are just participating to it, hence the “ING” name (without “P”).

Both ING and PING oscillations have been reproduced in computational models of networks of spiking neurons, where the firing of individual neurons and their synaptic coupling is modelled in detail (Whittington et al. 2000; Brunel, 2000). Neural mass models have also been designed, in which differential equations directly describe the properties of E and I populations involving many neurons (Deco et al., 2008). Most of these models are *phenomenological*, i.e. they embed stylized versions of physiological processes to emulate qualitatively the behavior of the original spiking networks. An example of such phenomenological models is the Rectified Linear Unit (ReLU) with delayed self-inhibition used in Battaglia, Brunel & Hansel (2007) to capture the collective dynamics of sparsely synchronized ING oscillations in spiking recurrent networks. Some models, however, are *exactly reduced*, since their macroscopic equations are rigorously derived from the ones describing the dynamics of the underlying spiking networks, modulo special assumptions on the neuronal dynamics and the properties of microscopic connectivity. For instance, the Montbrió-Pazó-Roxin model is exactly reduced from networks of Quadratic Integrate-and-Fire neurons (QIF) with quenched random excitabilities following a Lorentzian distribution (Montbrió et al., 2015). Such Exactly Reduced Models (ERMs) present the advantage of providing a precise mapping between microscopic physiological parameters and macroscopic effective constants. Combinations of Montbrió-Pazó-Roxin type dynamical equations describing motifs of coupled E and I populations have been able to reproduce both ING- and PING-type oscillations (Dumont & Gutkin, 2019). Here, to confirm the robustness of our theory and findings across alternative neural masses and oscillation-generation scenarios, we will focus on coupled Montbrió-Pazó-Roxin models to describe PING oscillations (Figure 1A) and on delayed Inhibition ReLU (dIReLU) models to describe ING-type oscillations (Figure 1B). Both these models have been previously shown to provide a faithful match (qualitative for the delayed-I ReLU, quantitative for coupled ERMs) between neural-mass and spiking simulations.

**Figure 1.**
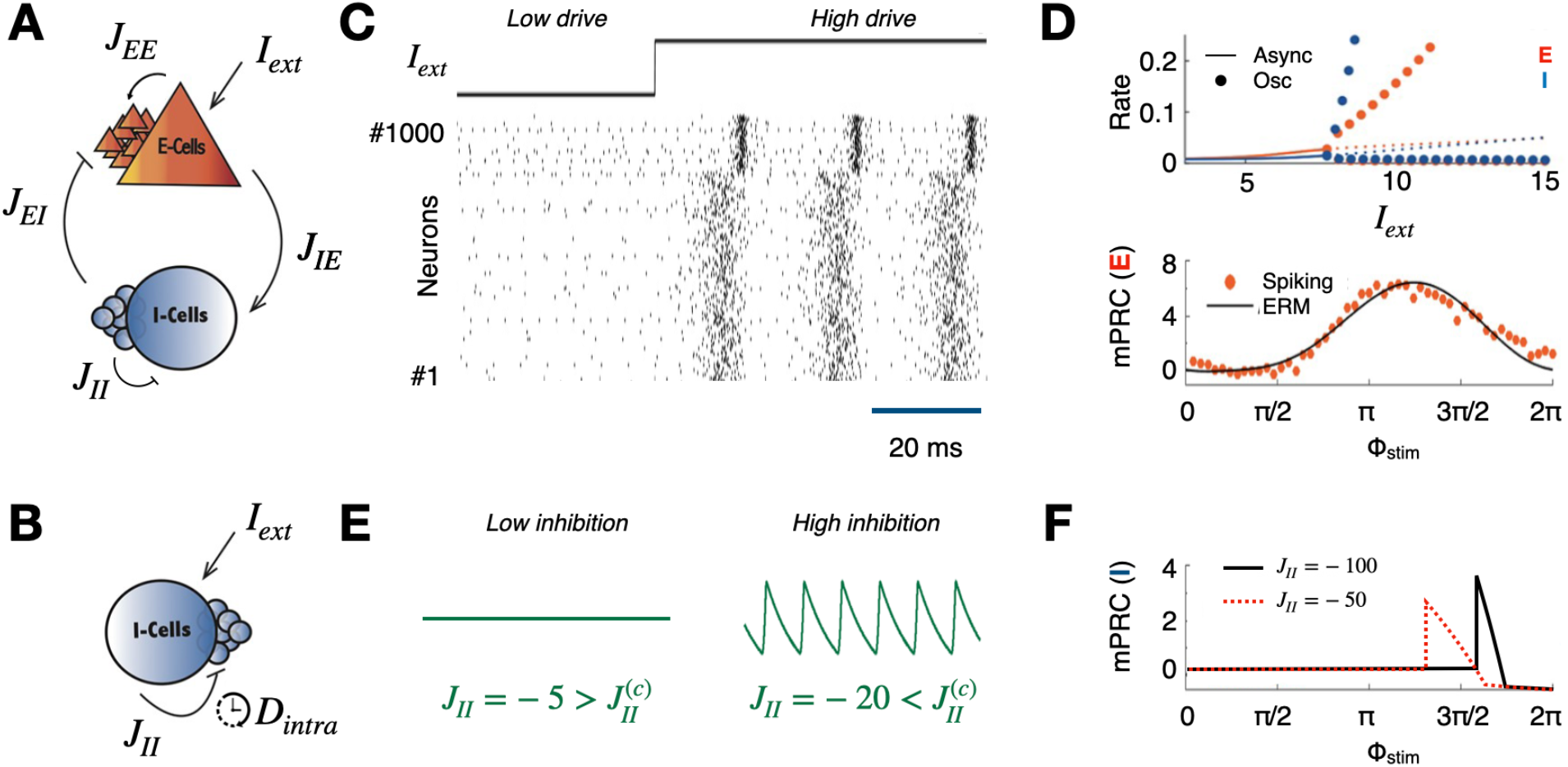
Models of regional dynamics. **A)** In the PING architecture (“pyramidal neuron / interneuron network generated” oscillations, each local region includes one excitatory (E) and one inhibitory (I) population coupled between them and with themselves through excitatory and inhibitory connections. **B)** In the ING architecture (“interneuron network generated”), each local region includes just one inhibitory population, coupled to itself with a delayed and negative inhibitory coupling. **C)** Example raster plot of a spiking network of Quadratic Integrate and Fire (QIF) neurons, undergoing a transition from asynchronous homogeneous to oscillatory firing as a function of the drive strength. **D)** These and other transitions of the QIF spiking network can be reproduced by a low-dimensional Exactly Reduced Model (ERM), for which we show here: a bifurcation diagram (top), where lines indicated stable fixed point and dots stable limit cycle solutions as a function of the drive *I*_*ext*_; a macroscopic Phase Response Curve (PRC, bottom), describing the (stimulation phase dependent) phase-shift of the local oscillation induced by applying a small intensity pulse perturbation at a given phase. Here the black line corresponds to a semi-analytic solution obtained via the adjoint method and the red dots to the result of numeric experiments of direct pulse stimulation on the spiking QIF network. **E)** Example rate time-series from a delayed Inhibition Rectified Linear Unit (dIReLU) implementation of the ING scenario. For weak inhibitory coupling *j*_*II*_, the activity has a constant firing rate, but for stronger inhibition 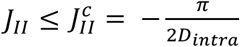 the rate begins to oscillate periodically. **F)** PRC curves of the ING diReLU model corresponding to different values of local recurrent inhibition strength.

In Figure 1C, we show an example simulation of a spiking network of QIF spiking neurons wired in a PING architecture as in Figure 1A. By varying the external drive, the spiking network undergoes a transition between an asynchronous regime of activity for low drive to a sparsely synchronized oscillatory regime for stronger drive. Such a transition can be predicted analytically, by analyzing a corresponding ERM (see Materials and Methods). As shown in Figure 1D (top), for low drive both E and I populations have a stable fixed point (solid lines), corresponding to constant average asynchronous firing rate. When the drive exceeds a critical current value, the fixed points become then unstable (dashed lines), while a stable limit cycle emerge (dotted lines), confirming the finding from spiking simulations. A similar transition occurs in the dIReLU implementation of the ING scenario (Figure 1B, see Materials and Methods) by changing the strength of the recurrent inhibition negative coupling *J*_*II*_, under the condition that inhibitory coupling is not instantaneous, but has a non-zero delay *D*. For weak local inhibition (Fig. 1E, left), the rate variable *R(t)* describing the activity of the unique ReLU node composing the model converge to a constant value, but, for stronger inhibition (specifically, *J*_*II*_ ≤ -π/2*D*, cf. Battaglia et al., 2007), *R(t)* can be proven analytically to undergo a bifurcation toward a limit cycle (Figure 1E, right), at a frequency decreasing for larger |*J*_*II*_|.

### Macroscopic Phase Response Curves

The analytical ERM and dIReLU models of PING and ING oscillations also allow predicting how the ongoing oscillations of the considered population would react to external perturbation inputs. As an effect of a brief but intense current injection, the ongoing oscillation could remain unperturbed or be delayed or advanced in phase. As measured empirically (e.g. Akam et al., 2012) and expected theoretically (Kuramoto, 1991), the phase-shifting effect of a pulse perturbation will depend on the phase of the ongoing oscillation at which the perturbation is applied: for some phases of application, the oscillations will be advanced in phase (positive Δφ); for others, it will be delayed (positive Δφ); for others yet, it will remain unperturbed (near-zero Δφ). This phase-dependency of the perturbation effect can be summarized by a function known as the *Phase Response Curve* (PRC) giving the induced phase-shift as a function of the phase of application of the pulse-perturbation (Pikovsky et al., 2003).

Note that we are considering here effects induced by the perturbation on the *collective* oscillations of the stimulated region. In other words, we consider perturbation effects at the macroscopic level of the entire region, not at the microscopic levels of effects on individual neurons. Individual cells in spiking networks indeed will usually have an irregular spike train even when the population collectively oscillate (Brunel, 2000) and cannot be considered individually as oscillators. For this reason, we speak of macroscopic Phase Response Curve, or mPRC (Dumont et al., 2017). In the PING ERM and the ING dIReLU models the mPRC functions can be evaluated analytically as a function of the model parameters. In Figure 1D (bottom) we show the mPRC for the excitatory population within the PING motif of Figure 1A, comparing direct measurements of phase-shifting effects of the applied perturbation in the spiking simulation (red dots) with analytical estimations from the corresponding ERM (well matching simulations). The shown mPRC indicate that pulse perturbations eventually advance the phase, with maximal effect for the second half of the oscillatory period. For more details on the mPRC estimation in the ERM, we refer the reader to the original publication (Dumont & Gutkin, 2019). Analogously, in Figure 1F, we show mPRCs for the ING dIReLU model for two values of the delayed self-inhibitory coupling *J*_*II*_. For this model, the collective oscillations is refractory to applied perturbations for broad phase intervals, as indicated by a zero-valued mPRC (see Battaglia et al., 2007 for details on the mPRC analytical derivation). Then, for other narrower perturbation phase ranges, the macroscopic phase can be both advanced or delayed.

As we will discuss in next section, the mPRC is also a key property for determining which phase-locking patterns between coupled oscillatory regions can be materialized in a stable manner.

### Both structural connectivity and regional dynamics affect inter-regional phase differences

The PRC of an oscillating local region describes its response to external perturbations. Now, the coupling to another oscillating region can be considered too as a perturbation. In the weakly-coupled phase oscillators theory, the phase shifting action exerted on the oscillations of region *X* by a connected remote region *Y* can be obtained by convolving the PRC of region *X* (ruling *X* phases responses to input pulses) with the input from region *Y* (conceivable as a continuous train of pulses of amplitude modulated by the oscillatory waveform of *Y*, scaled by the inter-regional coupling strength *G* and shifted by the inter-regional coupling delay *D*_inter_). At the same, time, in a system of two reciprocally coupled regions, the region *Y* will also see on its turn its phase further shifted by the input received from *X*. In short, a function Γ(Δφ) –written as the difference between two convolution integrals (Kuramoto, 1991; see Materials and Methods)– can be written giving the time-derivative of the phase-difference existing between the two coupled oscillating regions. Positive (or negative) values of such Γ(Δφ) will indicate that the current phase-difference is going to further dephase (or to reduce) over the next oscillation period. Of particular importance are thus the zeroes of this Γ(Δφ) function because they will indicate special phase difference values at which the two regions can *phase-lock* (in a stable manner if the slope of Γ(Δφ) at the zero point is negative, unstable otherwise). In the weak coupling limit, thus, knowledge of the Γ function is sufficient (and necessary) to determine which are possible phase-locking configurations of a system composed of two oscillating regions (and, hence, possible “functional connectivity motifs” since phase-difference are expected to modulate inter-regional communication, Bastos et al., 2014).

Noteworthy here is the fact that Γ(Δφ) does depend on the between-regions coupling parameters (strength and delay), as reasonable to expect. However, it also depends on the properties of the local oscillating regions, notably their PRC and oscillatory waveform, depending on their turn from other local parameters such as within-region inter-population couplings. The resulting phases of possible stable phase-locking will thus be difficult to predict based on knowledge of inter-regional connectivity alone. Eventually, inter-regional phase differences could be affected as much by alterations of local dynamics than by alterations of the inter-regional couplings themselves, as we now illustrate for simple systems of two symmetrically interconnected PING (Figure 2) or ING (Figure 3) regions.

**Figure 2.**
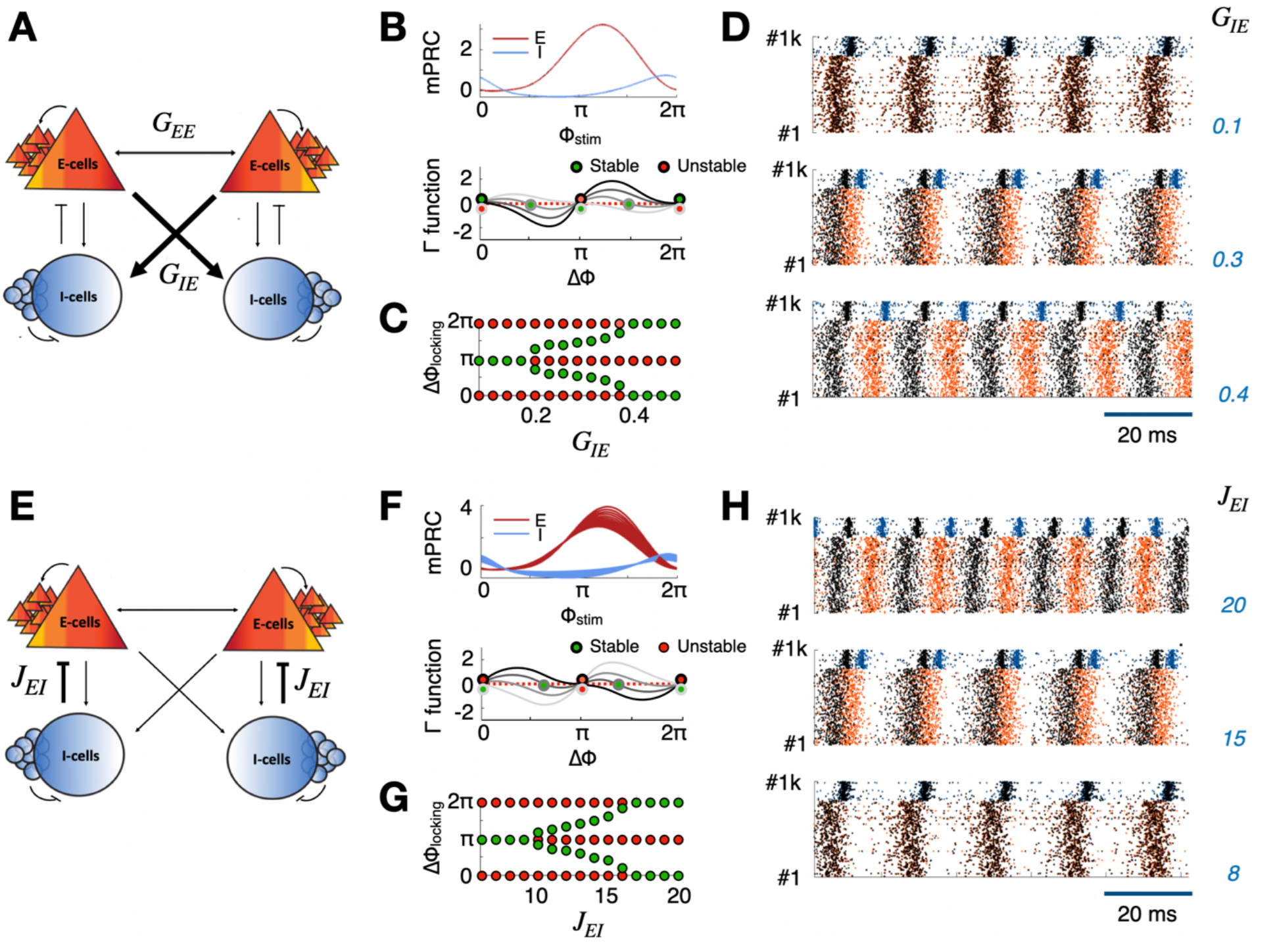
Phase-locking between coupled PING regions. Toy models of two bidirectionally and symmetrically coupled PING regions (as in Figure 1A) produce phase-locked oscillatory dynamics configurations. The obtained stable phase differences (in-, anti- or out-phase locking) depend on the properties of both the coupling and the local oscillatory dynamics. As an effect of modifying (**A)** inter-regional coupling strength (in this example the strength *G*_*IE*_ of the E-to-I interregional coupling, see Figure S1 for changes of the E-to-E inter-regional coupling *G*_*EE*_), **(B)** the Γ function ruling the dynamics of instantaneous inter-regional phase differences in the ERM model (bottom) is affected but not the local PRC of each population within the region. **C)** As an effect of *G*_*IE*_ decrease, the initial in-phase locking configuration is destabilized toward out-of-phase and, then, anti-phase locked configurations. **D)** A similar phenomenology is captured by simulating spiking networks of QIF neurons and reducing the strength of E-to-I long range synapses. The in-phase locking can however be restored by **(E)** reducing the I-to-E coupling *j*_*EI*_ within each of the regions. **(F)** Changes of *j*_*EI*_ affect the Γ function but now also the PRC of the local regions. **(H)** Once again, a similar phenomenology can be obtained in spiking networks of QIF neurons of which the low-dimensional neural mass model provides the exact reduction. See Figure S2 and S3 for the effects of changing, respectively, the local E-to-I coupling *J*_*IE*_ or the drive *I*_*ext*_ to the E populations within each region.

**Figure 3.**
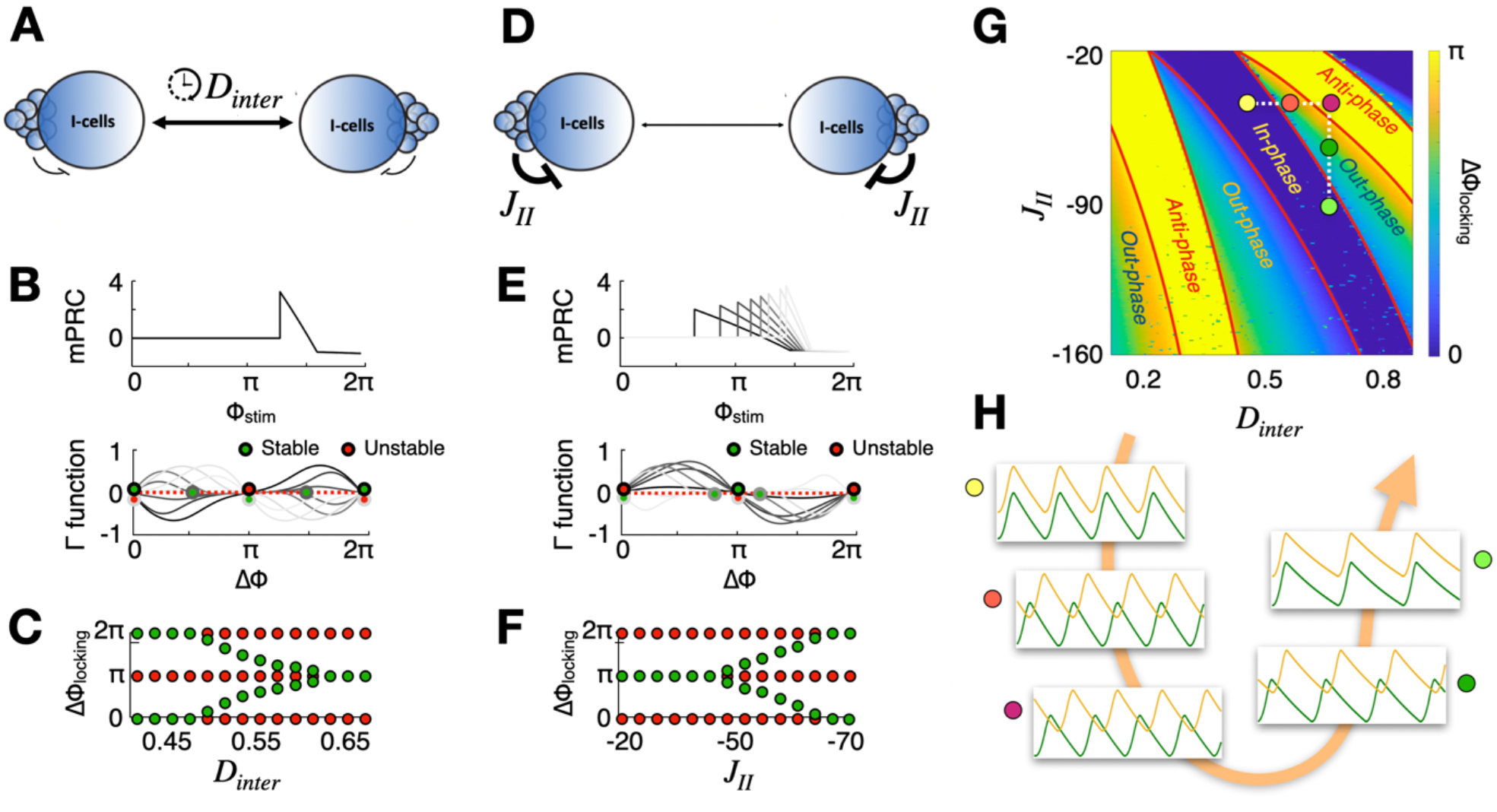
Phase-locking between coupled ING regions. Toy models of two bidirectionally and symmetrically coupled ING regions (as in Figure 1B) produce phase-locked oscillatory dynamics configurations. As for the coupled PING regions of Figure 2, the obtained stable phase differences (in-, anti- or out-phase locking) depend on the properties of both the coupling and the local oscillatory dynamics. **(A)** As an effect of modifying inter-regional delay *D*_*inter*_, **(B)** the Γ function ruling the dynamics of instantaneous inter-regional phase differences in the ERM model (bottom) is affected but not the local PRC of each population within the region. **C)** Increasing *D*_*inter*_, the initial in-phase locking configuration is destabilized toward out-of-phase and then anti-phase locked configurations. The in-phase locking can however be restored by **(D)** increasing the I-to-I coupling *j*_*II*_ within each of the regions. **(E)** Changes of *j*_*II*_ affect the Γ function but now also the PRC of the local regions. **(G)** Systematic analysis of the stable phase inter-regional differences obtained for changing *D*_*inter*_ and *j*_*II*_. Red lines correspond to semi-analytically determined boundaries of in- and anti-phase locking regions. Highlighted on the surface are six example configurations along a degradation (increasing *D*_*inter*_) and a compensation (increasing *j*_*II*_) pathways. **(H)** Time-series snippets from the highlighted points in panel **G**. Along the compensation pathway in-phase locking is recovered but oscillation frequency is slowed down. See Figure S4 for surfaces of amplitude and frequency of oscillation as a function of *D*_*inter*_ and *j*_*II*_.

We consider first the case of two mutually connected regions, each constituted by a PING system of two E and I populations (as in Figure 1A), modeled as ERMs (Figures 2B-C) or as actual spiking networks of QIF neurons. As shown in Figure 2A, these regions are coupled by excitatory connections orginating in the E population of each region and targeting both the E and I populations of the other region, with conductance strengths given respectively by *G*_*EE*_ and *G*_*IE*_ (in the ERMs) or changing probabilities of random connection (in the QIF networks). Figure 2B, shows how the PRC of each of the two identical regions and the Γ function vary as an effect of *decreasing G*_*IE*_ (see Supplementary Figures S1-S3 for variations of other parameters). The PRC of both the E and I populations (Figure 2B, top) depends uniquely on local coupling parameters, and, as such, is left unaltered by variations of inter-regional coupling strength. However, the Γ function is modified and notably changes its convexity, so that certain zeros can lose their stability and others can emerge along the variation of *G*_*IE*_ (Figure 2B, bottom). Figure 2C, tracks these changes of properties of zeroes as a function of *G*_*IE*_. For stronger coupling values, a stable in-phase locking configuration exists among the two populations. However, with the reduction of *G*_*IE*_, the in-phase configuration loses its stability and new out-of-phase locking configurations emerge, with either one of the populations leading in phase on the other. When the coupling is more strongly decreased, then a range of stable anti-phase phase-locking emerges. If we consider in-phase (anti-) as index of strong (weak) FC, then the degree of functional coupling between the two populations is weakened along the path of reduction of G_IE_. A similar effect can be achieved by reducing long-range E-to-I connection probability in spiking networks of QIF neurons, as visible in the sequence of raster plots of Figure 2D (see *Materials and Methods* for parameters). An opposite effect is obtained when varying G_EE_. An increase of G_EE_ destabilizes indeed the in-phase toward an anti-phase configuration, passing through out-of-phase locking patterns (cf. Figure S1).

We can then modify the phase-locking configuration of the PING bioregional network by acting on local dynamics within both regions. Figure 2E-H refer to an *in silico* experiment in which we manipulate the intra-regional inhibition to excitatory neurons coupling *J*_*EI*_. Such manipulation perturbs the internal balance between E and I currents received by each population and, as such, modulates the excitability of the considered regions. As computed analytically for the ERM, under a variation of local dynamics induced by changes in *J*_*EI*_ both the PRCs and the Γ function are now affected (Figure 2F). As a result, the considered configuration of anti-phase locking between the two populations achieved by reducing G_IE_ in Figures 2A-D is restored into a stable in-phase locking pattern (Figure 2G). Once again, a similar effect holds for the spiking QIF network model (Figure 2H).

An opposite effect is obtained by modulating *J*_*IE*_ (Figure S2) or the external current drive to the excitatory population (Figure S3).

We consider then the case of two coupled ING regions, of the type of Figure 1B. In this case each region contains just an inhibitory population, self-inhibiting itself with a delayed coupling, with a delay *D*_local_. We model here regions through a dIReLU neural mass, even if analogous spiking simulations could be constructed (cf. Battaglia et al., 2007). The coupling between the two populations is anyway of excitatory nature, even if no explicit E population is modeled in this case. We first perturb this coupling by modifying its speed of signal propagation, resulting in an increased inter-regional delay *D*_inter_ (Figure 3A). As for the case of the PING motif, modifying coupling parameters leave unaffected PRCs but alter the Γ function (Figure 2B). Here, specifically, increasing *D*_inter_ destabilized an in-phase locking configuration into an anti-phase, passing through antiphase (Figure 3C, a similar effect to changing *G*_*IE*_ in the PING motif, cf. Figure 2B). The reached phase-locking configuration can however be restored into an again stable in-phase locking, by increasing in both regions the local recurrent inhibition strength *J*_II_ (Figure 3D-F, similar to changes of *J*_EI_ Figures 2E-H).

For this ING bioregional motif, we show in Figure 3G the overall dependency of inter-regional stable phase-difference as a function of changing long-range delay (a connectivity parameter) or local inhibition (a regional parameter). For every given value of *D*_inter_, every possible inter-regional phase-relation can be stabilized by choosing a suitable value of *J*_II_. This indicates very transparently that variations of inter-regional phase difference and functional coupling cannot be straightforwardly attributed to variations of the underlying structural coupling. Indeed, this phase-relations are determined through a complex interplay between coupling and regional dynamics properties, allowing variations of local dynamics to compensate for the phase-difference variations induced by structural changes.

An example of this compensation can be obtained by following the dotted lines highlighted on top of Figure G and corresponding to the parameter changes occurring through the concatenated steps of Figures 3A-C (*“degeneration path”*, as structural coupling is getting slower, as in a “toy” demyelinating neurodegenerative disease) and Figures 3D-F (*“compensation path”*, as the structural coupling remains degraded but an additional modulation of local dynamics compensates for the effects of this structural degeneration). Along these paths, the in-phase synchrony associated to strong FC is first decreased then reinstated. The final, compensated condition is, however, not completely identical to the starting one. As visible in the time-series snippets shown in Figure 2H for selected points on the paths of Figure 3G, the in-phase oscillations of the compensated point have a reduced frequency with respect to the original condition, prior to coupling degeneration. We measure here time in arbitrary units, however, if we considered the frequency of the oscillation at the starting yellow point to be, conventionally, ∼30 Hz, then the frequency of the final light green point along the path would be of ∼23 Hz, i.e. a ∼20% relative variation. Variations of frequency as a function of *D*_inter_ and *J*_II_ can be seen with greater detail in Supporting Figure S4A, together with the (milder) variations of oscillatory amplitude in Figure S4B.

### Degeneration and compensation paths in large-scale whole-brain models

We have so far shown examples of SC-degeneration induced FC changes, later compensated by local dynamical changes, only in toy models involving two symmetrically connected regions. However, brain networks are complex systems composed of much larger number of regions, coupled by heterogeneous connectivity, possibly asymmetric. To probe the robustness of the compensation by dynamics mechanisms probed in Figures 2 and 3, we then study their feasibility in large-scale whole-brain connectome-based models. In this models, neural mass systems describing regional dynamics are attached at each node of a multi-regional SC connectivity matrix derived from empirical reconstructions (Honey et al., 2007; Deco et al., 2011). The spontaneous, noise-driven dynamics of these systems is then simulated, generating time-series out of which matrices of correlation-based FC can be estimated and compared to equivalent empirical data, if available.

Here we consider two example model architectures. In Figure 4, we adopt a generic human connectome matrix mediated from Hagmann et al. (2008) and previously used in many simulation studies (e.g. Deco et al., 2013; Hansen et al., 2015). For such connectome matrix, shown in Figure 4A, tract-length information providing indication of delays of propagation is also available and we chose therefore to use it in conjunction with an ING dIReLU regional dynamics to test the feasibility of *delay-increase compensation* as in Figure 3. Such SC matrix is obtained via Diffusion Tensor Imaging and is therefore heterogenous, but still symmetric as in the toy models. To test, therefore, conditions even farther away the one of the toy models, we consider in Figure 5, an asymmetric and highly heterogeneous SC matrix (Figure 5A) obtained via direct tracer injection experiments in non-human primate brains (Markov et al., 2014). We use then this matrix in combination with PING ERM models of regional dynamics to test the feasibility of *connectivity-loss compensation* as in Figure 2.

**Figure 4.**
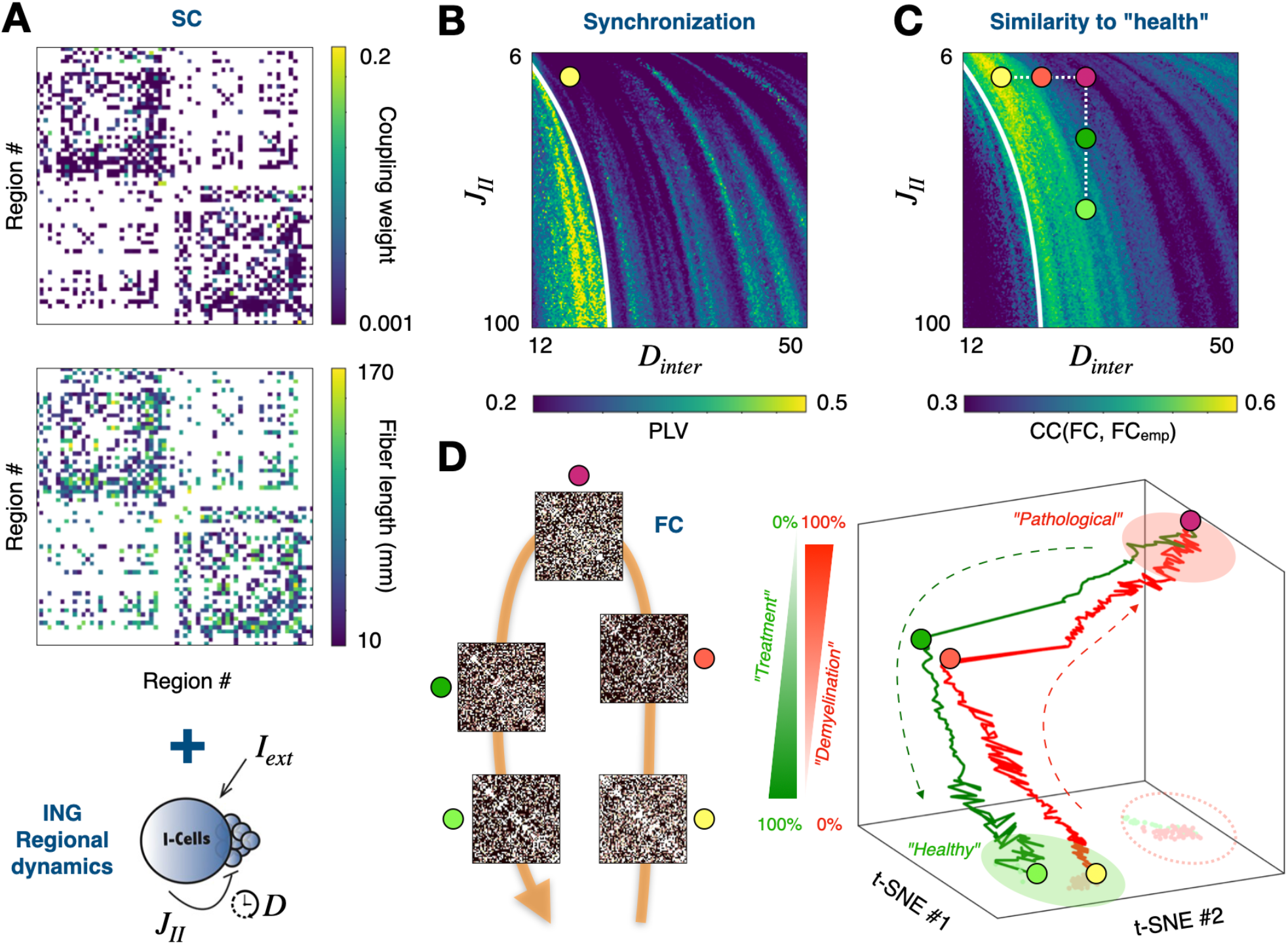
Compensation of degeneration in a whole-brain model with ING dynamics. **A)** Human DSI-derived connectome strength (top) and length (bottom) used in combination with ING dynamics for large-scale simulation. A “healthy” Dynamic Working Point of operation (in yellow) is selected **(B)** in proximity to a transition to a regime enhanced inter-regional synchronization (tracked by average Phase Locking Value, PLV), in a point which precisely maximize **(C)** the similarity between the simulated FC matrix and an empirical FC_emp_ measured from fMRI in healthy subjects. Such similarity is maximized in a regime slightly subcritical with respect to the synchronization transition. Simulated demyelination (increase of *D*_*inter*_) causes the system to leave this regime along a degeneration pathway. However, increasing *j*_*II*_ within each region causes the system to follow a compensation pathway entering again a zone of higher similarity to healthy FC_emp_. **(D)** To the left are shown FC matrices obtained at the points along degradation and compensation pathways highlighted in panel **C**. Also shown to the right is a dimensionally reduced representation of FC matrix changes along these pathways, obtained via a non-linear distance preserving t-SNE projection of the sequences of simulated matrices (the vertical dimension reports % progression along the degeneration and compensation pathways, the projections of the “healthy” and “pathological” point clouds on the tSNE 2D latent space are denoted by green and red ellipses).

**Figure 5.**
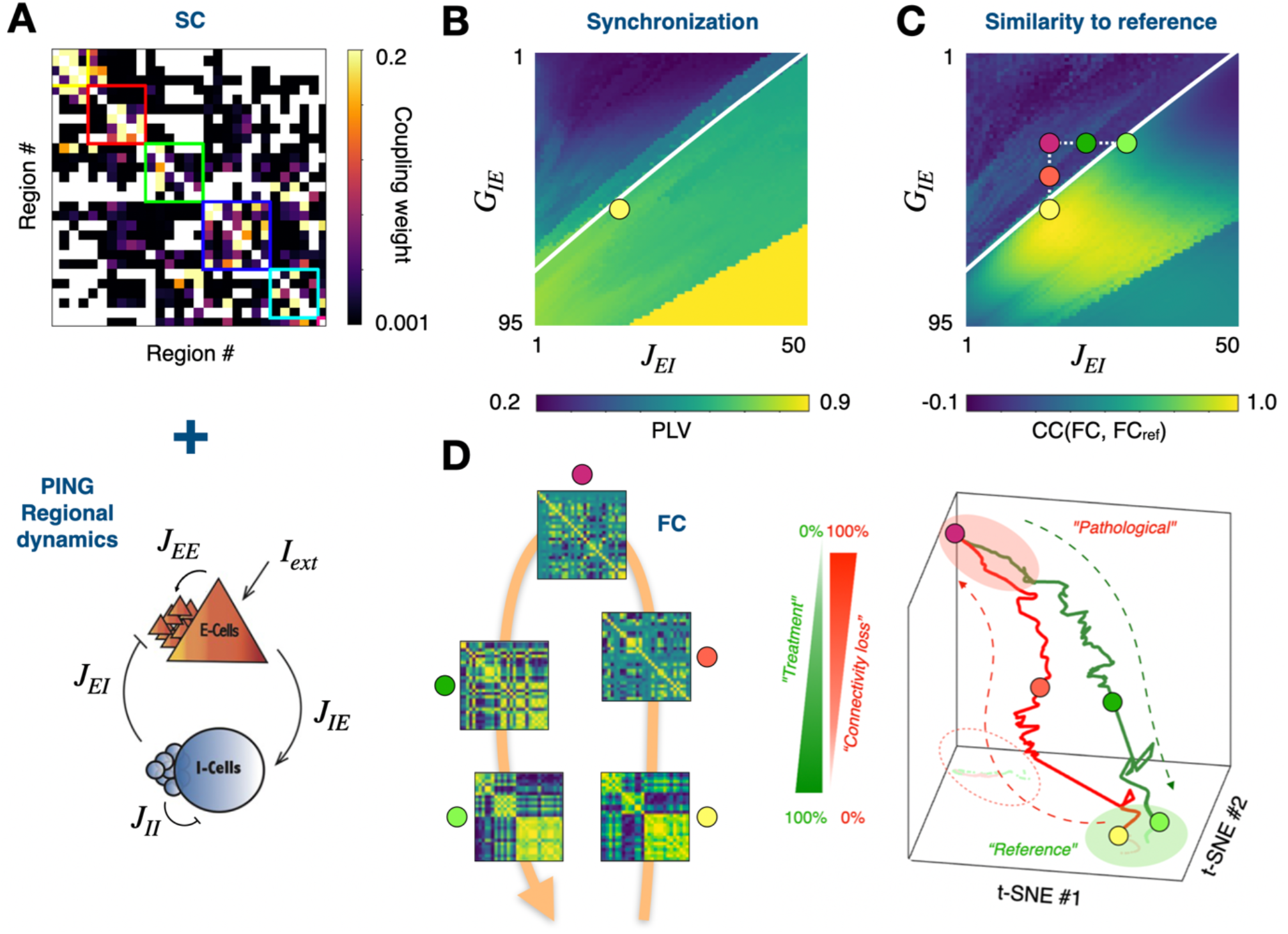
Compensation of degeneration in a whole-brain model with PING dynamics. **A)** Non-Human-Primate connectome strength matrix derived from tracer injection experiments used in combination with PING dynamics for large-scale simulation. **(B)** A “reference” Dynamic Working Point of operation (in yellow) is selected in a regime with mixed inter-regional synchronization (as tracked by average Phase Locking Value, PLV), close to a critical transition (white line) from a desynchronized state. (**C**) Similarity between the obtained FC and the reference FC_ref_ matrix as a function of changing DWP. Simulated connectivity loss (decrease of *G*_*IE*_) causes the system to leave this regime along a degeneration pathway. However, increasing *j*_*EI*_ within each region (as in Figure 2E) causes the system to follow a compensation pathway entering again a zone of higher global PLV. **(D)** To the left are shown FC matrices obtained at the points along degradation and compensation pathways highlighted in panel **C**. Also shown to the right is a dimensionally reduced representation of FC matrix changes along these pathways, obtained via a non-linear distance preserving t-SNE projection of the sequences of simulated matrices (the vertical dimension reports % progression along the degeneration and compensation pathways, the projections of the “reference” and “pathological” point clouds on the tSNE 2D latent space are denoted by green and red ellipses).

The first step in tuning a whole-brain connectome-based model consists in the identification of its Dynamic Working Point (DWP), determined by a choice of global parameters, here a scale *G*_*inter*_ of strength and, optionally, a delay scale *D*_*inter*_. Strengths and latencies of individual pairwise connections are then obtained by multiplying these scales by the connectome strength and connectome length empirically reconstructed matrices. Different choices of DWP lead to different dynamical regimes and, thus, to different resulting FCs. Some criteria are then required to choose the DWP. In some cases, an empirically determined FC matrix is available. Therefore, the DWP can be determined by maximizing the correspondence between the simulated and empirical FCs. This is the procedure which we follow in Figure 4 for the human symmetric connectome with ING dynamics model, since an empirical FC companion to the adopted SC is available (cf. optimizations of DWP in Deco et al., 2013; or Hansen et al., 2015). The optimal DWP thus determined is reported in *Materials and Methods* and graphically represented by a yellow dot in Figure 4B. In some other cases, a companion FC is not available. Yet, a plausible DWP can be guessed based on the qualitative properties of the dynamics associated with it. For the directed connectome of Figure 5A, used in combination with a PING dynamics, we don’t dispose of matching FC information. We follow therefore Benitez-Stulz et al. (2023) and select a DWP nearby a critical line between a phase un-locked regime and a heterogeneously phase-locked regime, conferring rich dynamics with the coexistence between a core of synchronized regions paired to a residual dynamics of FC (in line with general recommendations on DWP selection discussed by Arbabyazd et al., 2021).

Starting from the analysis of the ING whole-brain model on a human connectome (Figure 4), we observe that, for the chosen value of *G*_*inter*_, a critical line exists in the parameter plane spanned by the inter-regional delay *D*_*inter*_ and the regional self-inhibition *J*_*II*_, separating a region of higher from a region of lower inter-regional synchronization (as measured by variations of the PLV index, see Materials and Methods and Figure 4B). In Figure 4C, we show the variations on the same parameter plane of the similarity of the obtained simulated FC with a reference empirically determined FC. This similarity is larger in a slightly subcritical stripe bordering the critical transition line to a high synchronization regime and is specifically maximized for the chose DWP, as previously mentioned. Attempting to reproduce in the large-scale model the degeneration and compensation pathways of Figure 3G, we first increase the inter-regional delay *D*_*inter*_ to emulate phenomenologically the effects of progressive demyelination. As visible in Figure 4C, this degeneration path cause the DWP to shifts away from the slightly subcritical stripe and, consequently, the simulated FC gets perturbed and increasingly deviating from the “healthy” baseline condition given by the FC at original DWP. Once again following Figure 3G, we then increase self-inhibition *J*_*II*_ unspecifically in all regions (see *Discussion* for potential mechanisms that could achieve this effect: medication, changes of neuromodulation…). Following the vertical compensation path, the DWP enters again the slightly subcritical stripe giving FCs bearing similarity the healthy empirical FC conditions, leading to a “repaired” FC closer to the original one, despite the inter-regional delay remaining increased. The degeneration of SC is thus still existing, but the same dynamical regime can still be accessed by modifying local dynamics. Figure 4D (left) shows example simulated FC matrices corresponding to the marked dots along the paths of Figure 4C. Yet another representation of the degeneration path and of the return closer to healthy normality is presented in Figure 4D (right). Here, we make use of a non-linear distance-preserving dimensional reduction technique (t-stochastic neighborhood embedding, Hinton & van der Maaten, 2008) to project in a bidimensional space (spanned by two abstract latent variables tSNE #1 and #2) the matrices of FC generated along the degeneration and compensation paths. These FC matrices and the paths connecting them are thus represented as points and lines in the latent space (degeneration path in red, compensation path in green). Here, to better visualize the progression along the paths, we add an additional third vertical dimension, indicating the progression along the paths, so that the last point along the degeneration path has the largest vertical elevation and the last point of the compensation path is again at zero vertical elevation. Red and green zones reported on the bidimensional latent space indicate a zone surrounding the end points of the degeneration and compensation paths, for the sake of a simpler comparison. We observe that the degeneration path brings FC in a region far from the initial one, but the termination of the compensation path falls again in a closer neighborhood of the healthy FC, prior degeneration. We thus achieve a straightforward visualization of FC compensation, despite the large dimensionality of the simulated multi-regional system.

Figure 5 shows an analogous construction of degeneration and compensation pathways for a PING model on top of the directed NHP connectome of Figure 5A. The selected DWP –the one that we conventionally choose to associate to a “healthy” references in this case– lie at the edge of a critical phase unlocked/locked transition in the parameter plane spanned by the inter-regional coupling strength *G*_*IE*_ and the regional I-to-E coupling *J*_*EI*_. (Figure 5B). Once again, by reducing *G*_*IE*_ as in Figure 2B, to phenomenologically emulate weakening of connectivity fibers, the DWP of the system leaves this edge-of-synchrony regime to enter a more desynchronized regime. However, the increase of *J*_*IE*_ causes it to enter again into a more synchronized regime, despite the connectivity loss not being restored (Figure 5C). Analogously to Figure 4D, we show in Figure 5D example FC matrices along the degeneration and compensation pathways, as well as t-SNE projections of these same pathways, so to clearly depict: the divergence from reference conditions caused by degeneration; and the return to the neighborood of them thanks to compensation.

As in the case of the PING toy models of Figures S1, S2 and S3, additional degeneration and compensation pathways can be considered. In Figure S5, we compensate the decrease in *G*_*IE*_ in the PING connectome-based model by raising the value of the regional background drive *I*_*ext*_ (cf. Figure S3). In Figure S6, we consider rather than degeneration of the connectome a plastic change in which the strength of *G*_*EE*_ increases (cf. Figure S1). However, even in this case the original FC can be preserved –implementing a form of “FC homeostasis” (Marder & Goaillard, 2006; Kisperky et al., 2011) – by increasing the local couplings *J*_*IE*_ (cf. Figure S2). In these additional examples we also use modified DWPs (e.g. slightly supercritical in Figure S5 and precisely critical in Figure S6, rather than slightly subcritical as in Figure 5) to show that compensating global FC changes by local dynamics modulations can be robustly implemented for different starting reference conditions.

In conclusion, the compensation phenomena identified for the ING and PING toy-models in Figures 2 and 3 (and S1–S3), are reproduced as well when these models are embedded into much more complex multi-regional networks with realistic connectomes in Figures 4 and 5 (and S4–S5). This allows to hypothesize that compensation of SC damage by dynamics modification could be contributing to “cognitive reserve” in neurodegenerative diseases and FC homeostasis across brain plastic changes (see *Discussion*).

## DISCUSSION

Structural and Functional Connectivity (SC and FC) are inter-related but not completely redundant, as FC is the manifestation of dynamics unrolling on top of SC and dynamic complexity can produce some “special effects”. One of them is the possibility of functional multiplicity (many FCs stemming out of a same SC, Battaglia & Brovelli, 2020). Here we focus on the less frequently explored, complementary phenomenon of structural degeneracy, in which similar FC states can arise from diverse SCs, if they produce a shared emergent dynamical regime. Structural degeneracy arises indeed in the ING and PIN toy models of Figures 2 and 3 and in an even more striking manner in the large-scale connectome-based models of Figures 4 and 5. Matching phase-locking configurations associated to efficient in-phase synchronization or similar matrices of FC can be obtained starting from diverse SC connectomes with increasing degrees of degeneration (decreased *G*_*inter*_ or increased *D*_*inter*_, mimicking respectively loss of connectivity or demyelination). Indeed, suitably tuned variations of local dynamics can precisely compensate for the detrimental effects of degeneration on FC, maintaining the system in the same global dynamical regime as before SC was damaged.

This structural degeneracy at the meso- and macro-scale of multi-regional networks is not dissimilar from “functional homeostasis” observed in invertebrate micro-scale circuits, where very different synaptic parameters and neuronal conductance values lead to strongly preserved movement patterns (Marder & Goaillard, 2006). In the case of invertebrate central pattern generators, however, differences between the underlying structural circuits, concealed by the generation of a common target dynamics in normal operating ranges, are suddenly revealed in the case when external variables as temperature change in an environmental shock (Tang et al., 2012). Here, the target FC state is associated to well-delimited zones in the parameter space conferring the desired properties, e.g. in phase locking in Figure 3G, similarity to empirical healthy FC in Figure 4C or yet elevated collective synchrony in Figure 5B. The boundaries of this zone may sometimes be given by critical transition lines in system’s dynamics (e.g. a sync/desync transition), so that we predict a gradually progressing degeneration may have little effects on FC until when one of these critical transition lines is trespassed. This applies also to the compensatory paths, where the progress toward a restored FC may be initially slow but displays sudden improvements, while control parameters of local dynamics are continuously changed (some abrupt changes are visible for instance in both the degeneration and the compensation paths in Figure 4D). Crossing of critical transition lines delimiting dynamical regimes and constituting thus boundaries of sudden FC change may speculatively provide an explanation for fast fluctuations of cognitive performance sometimes observed in patients of neurodegenerative diseases (Palop et al., 2006).

In this “functional homeostasis” view, the resilience of cognitive efficiency in a subject affected by neurodegenerative processes or even just undergoing healthy aging (Snowdon, 2003; Rentz et al., 2010), a capacity usually attributed to cognitive reserve processes not yet fully elucidated at the circuit level (Stern et al., 2003; Lee et al., 2019), could be enabled by fine dynamics tuning mechanisms that keep the system within the boundary of a “healthy” or “health-like” state of collective dynamics. For compensation via dynamics to occur it would not be necessary to drive the system in exactly the same Dynamic Working Point (DWP) as prior to SC degradation but steering it toward an alternative DWP lying within the same dynamical regime could be sufficient. This is what happens in the conceptual simulated experiments of both Figures 4 and 5. In Figure 4, for instance, the final DWP reached after compensation engenders a FC close to the original, pre-degeneration FC, but the oscillatory frequency of regional oscillations is, in this new DWP, reduced. Reduced frequency of oscillations are often observed in aging or pathologies (Osipova et al., 2005; Insel et al., 2012; Wiesman et al., 2021) and sometimes interpreted negatively as a manifestation of reduced speed of processing (Salthouse, 2000). However, our study suggest that these frequency modulations could also be the epiphenomenal signature of positive compensatory mechanisms at play.

One mechanism to potentially implement the required changes of synaptic couplings *J*_*II*_ or *J*_*EI*_ could be fine neuromodulatory adjustments, that can induce widespread changes of local synaptic efficacies (Brunel & Wang, 2001). The well-documented dysfunctions of neuromodulatory systems in neurodegenerative diseases (Martorana et al., 2009; Nobili et al., 2017; Hampel et al., 2019) could thus contribute to symptoms worsening via a less effective implementation of compensation of degeneration by the fine-tuning of dynamics. Note that neuromodulatory adjustments may also support the homeostatic maintainance of specific FC networks under white matter tracts plasticity (Sampaio-Baptista & Johansen-Berg., 2017) which may increase net long-range coupling efficiencies (as in Figure S6) rather than degrading them.

Control and adjustment of non-local FC interactions could also be achieved via the interplay of multiple inhibitory interneuronal classes within each population, which provide precise circuit-level mechanisms for the fine regulation of the overall level of inhibition (Burkhalter, 2008). Indeed, as predicted by the analyses of Figure 3G, adjustments of local inhibition may provide a powerful knob for the shifting of inter-regional phase differences. Rich longitudinal studies, with systematic assessment of connectivity degeneration, neurotransmitter and neuromodulator levels as well as precise cognitive profiling in addition to FC analyses will be ultimately needed to probe the pertinence of our hypotheses.

If our theory was correct, it could become conceivable to design treatments that do not aim at reinstating the function of individual system’s components but directly the collective dynamical regime of the whole system. Global and not specific adjustments of synaptic conductances needed to produce emergent compensation by dynamics could be obtained through pharmacological drugs that have direct modulatory effects on the strength of GABAergic transmission, such as Benzodiazepines. Note indeed that altered cerebrospinal fluid levels of GABA are indeed observed in Alzheimer’s disease and benzodiazepine treatments are sometimes prescribed, although with mixed results (Salzman, 2020). The heterogeneity across patients of the effects of these treatments may be explained by the need of carefully tune the administered dosage to steer brain networks in the appropriate regime. In the future, therefore, the model-driven personalized optimization of drug dosages could become a design solution of primary importance. Finally, in Figures 2 to 5 we focus on obtaining compensatory changes of dynamics by acting on synaptic conductances. However, Figures S3 and S5 predict that inter-regional phase differences could be modified also by acting on the drive currents to the coupled populations. The use of stimulation could then also be envisaged for FC control. However, the spatio-temporal patterns of applied stimulation should be carefully designed accounting for both the target FC state to achieve and the current FC state into which stimulation is applied, as effects are expected to be nonlinear and region-, phase- and state-dependent (Battaglia et al., 2012; Kirst et al., 2016; Benitez Stulz, 2023).

For both stimulation design and drug protocol optimization, simulation-based approaches are expected to play a growingly important role, given their capacity to account for and reproduce dynamic complexity. In this context, the use of Exactly Reduced Models for describing local dynamics –as the PING ERM of Figures 2 and 5– is particularly adapted. Indeed, the parameters appearing in the mesoscopic dynamic equations of ERMs are directly interpretable in terms of the underlying spiking network details, such as densities and strengths of synaptic connections, which are the microscopic level substrate on which any medical drug will concretely act (even if what treatments attempt to trigger are changes of meso- and macroscopic dynamics). Here we strongly focused on FC changes obtained by reconfigurations of oscillatory dynamics –a prominent driver of FC flexibility (Fries, 2005)– and emphasized therefore the role of PRCs in the analytical understanding of how circuit parameters affect phase differences (Figures 2 and 3). The PRCs we considered directly describe the phase response of collective oscillations of neuronal populations and are thus macroscopic FCs (Dumont et al., 2017), not be confused with microscopic PRCs which describe the phase response of individual neurons within the population. Importantly, such macroscopic PRCs can be analytically computed for models such as the ING diReLU and the PING ERM we here considered. They could however be measured in computational simulations for more complex neural mass models whenever they were needed. Note that the PRC framework has been extended to encompass the case of amplitude fluctuations (Castejón et al., 2013) or of multiple frequencies interacting (Ceni et al., 2020) as well. These developments could then be used in future studies exploring the robustness of compensation by dynamics mechanisms in more complex scenarios than the regular, single frequency situations here explored. Moreover, ERMs and neural masses are not limited to the reproduction of oscillatory dynamics and also account for other dynamical phenomena such as rate multi-stability (Wong & Wang, 2006) which also may shape collective dynamics and thus FC (Deco et al., 2013; Hansen et al., 2015). Therefore, we expect that the concept of dynamic compensation –particularly transparent in the systems with oscillatory dynamics considered in this study– is pertinent even for more general, non-oscillatory dynamical regimes.

The toy and large-scale models here considered were very abstract and still unfit to describe actual neurodegenerative diseases, where connectivity loss and demyelination damages are spatially heterogeneous and have very specific paths of progression (Braak & Braak, 1991). Yet, our models provide a first proof of concept of the potential impact on FC of compensation by dynamics phenomena, usually neglected in structure-centered views of SC-to-FC interrelations. Databases of patient-specific, disease-specific alterations of SC are increasingly available –see e.g. the FTRACT database for precise and systematic evaluation of inter-regional couplings directionality and latency (Trebaul et al., 2018)– and could be used to constrain pathology and patient-specific models to be used in real personalized applications.

Waiting for these pragmatic improvements, our general models allow since now advancing in our theoretical understanding of FC control. Specifically, they open interesting perspectives about an important overarching question: what is the ‘cause’ of functional deficits in cognition and behavior arising through pathology or aging? Here a mock neurodegeneration –representing connectivity loss or demyelination– induces alterations of FC, a proxy for functional impairments in real subjects and patients. However, compensation by dynamics allows restoring this FC without need to correct for the occurred SC degradation. This means that what we are “repairing” is not the connectome but the dynamical state, responsible for FC emergence more than the underlying SC. In this view, the ‘cause’ of the functional impairment would be algorithmic rather than structural (Marr & Poggio, 1976), i.e. identified into the change of system’s dynamics itself more than into the specific neurodegenerative process that led to it (which is just a mechanism to obtain a dynamical change, but not exclusive, and thus possible to counteract). According to our views, neurodegenerative diseases should thus be considered as “dynamopathies”, like epilepsy, more seriously than usually thought (Petkoski et al., 2023).

## MODEL AND METHODS

### Quadratic-Integrate and Fire spiking networks and their exact reduction

The reader is referred to Dumont & Gutkin (2019) for full details and model derivation. We provide nevertheless here some summary information. The model is obtained from the reduction of an an all-to-all coupled network made up of *N* spiking cells characterized by the quadratic integrate-and-fire (QIF) model:

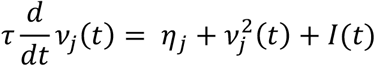

where *v*(*t*) represents the time evolution of the membrane potential, *τ* is the membrane time constant, *I*(*t*) is the total current, and we assume the intrinsic parameter *η* being randomly distributed across the network according to a Lorentzian distribution :

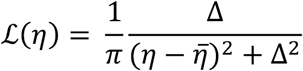

with mean value 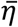 and distribution half width Δ. Such parameter emulated the variability of excitability induced by quenched disorder. The onset of an action potential is taken into account by a discontinuous mechanism with a threshold *v*_*th*_ and a reset parameter *v*_*r*_ respectively set at plus and minus infinity. Spiking simulations of QIF networks shown in Figures 2, S1-S3 are performed by direct integration of the QIF equations, using *N*_*E*_ = 4000; *N*_*I*_ = 1000 neurons (*v*_*threshold*_ = 500; *v*_*reset*_ = *−*500.

A mean-field limit analysis of the population firing rate 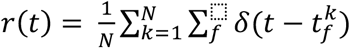, where 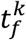 are the times of the *f-*th spike emitted by neuron *k*, has been conducted by Monbrió, Pazó and Roxin (2015), once again exploiting a Lorentzian ansatz for the probability *p*(*t, v* |*η*) of observing any randomly chosen cell with a potential *v* at time *t*, knowing its intrinsic excitability parameter *η*. After various algebraic manipulations under these assumptions, an equation for *r(t)* can be explicitly derived:

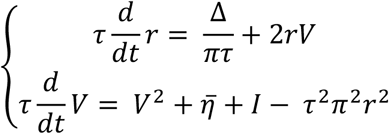

where V is the Lorentzian-averaged membrane potential of *v* over the population.

### Exactly Reduced Models of (coupled) PING regions

To construct a PING region, following again Dumont & Gutkin (2019), we couple the ERMs for two distinct E and I populations:

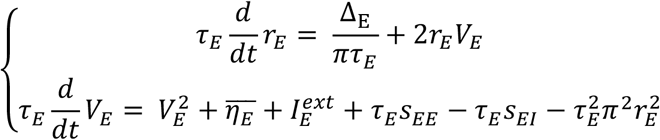

and:

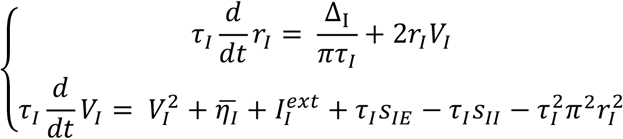

Here, the terms *s*_*αβ*_*(t)* are synaptic currents sent from population β to population α with α, β = E or I. These synaptic currents have exponential decay after each spike and evolve with dynamic equations:

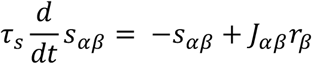

where *τ*_*s*_ are synaptic time constants and *j*_*αβ*_ synaptic strengths. For PING region simulations in the article we use the following values: *τ*_*E*_ = *τ*_*I*_=10 ; 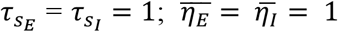; *j*_*EE*_ = *j*_*II*_ = 0; *J*_*IE*_ = *j*_*EI*_ = 15 (when not otherwise specified); 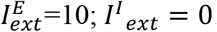. When coupling together multiple regions, every region includes a separate E and I population. The E_pre_ population in each presynaptic region sends connections not only to the I population within the region itself but also to the E and the I populations in each remote population. These inter-regional synaptic currents follow the same equation as for intra-regional currents:

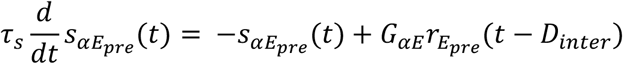

The presynaptic E population rate appears in the differential equation for inter-regional currents, scaled by two conductance multipliers *G*_*EE*_ (pre-synaptic to post-synaptic E) and *G*_*IE*_ (presynaptic E to postsynaptic I), and delayed by the delay of the inter-regional connection *D*_*inter*_ In the two PING regions toy model of Figure 2, we use *G*_*EE*_ = 0.1; *G*_*IE*_ = 0.5; *D*_*inter*_ = 6 (when not otherwise specified).

### Exactly Reduced Models of (coupled) ING regions

As a model of local ING-generating region, we use a delayed Inhibition Rectified Linear Unit model (dIReLU), introduced by Battaglia, Brunel and Hansel (2007). In this model every region includes just a single inhibitory population coupled to itself by a delayed self-inhibitory loop:

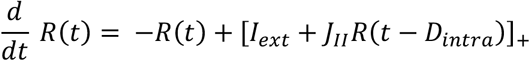

where [x]_;_ = x if x ≥ 0, or 0 otehrwise and *D*_*intra*_ (or shortly *D*) is the local delay within each region and *j*_*II*_ *<* 0, denoting dominantly inhibitory local recurrent interactions.

When coupling two regions, an additional long-range inter-regional coupling is added:

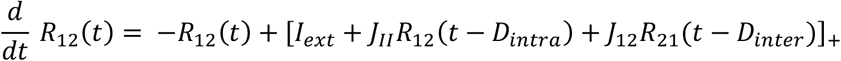

Where *j*_1*2*_ > 0 corresponds to an excitatory influence of region *2* (or 1) to region 1 (or *2*). In general, *D*_*inter*_ ≥ *D*_*intra*_ Here, we use *j*_1*2*_ = *j*_*2*1_ = 1 and *D*_*intra*_ = 0.1. Values of *j*_*II*_ and of *D*_*inter*_ are specified for every analysis in the figure caption.

### Large-scale connectome-based models

The equations for the large-scale models of Figures 4 and 5 are straightforward generalizations of the previously described dynamic equations of ING and PING models, respectively. Specifically, for the large-scale ING model of Figure 4, coupling conductances between regions *i* and *j*, proportional to an adjustable scale of long-range excitation *j*_*inter*_, are further scaled by the relative connectome strength *g*_*ij*_ given by the adopted connectome matrix, and inter-regional delays by the tract lengths Λ_*ij*_. The dynamic equation of a region *I* thus becomes:

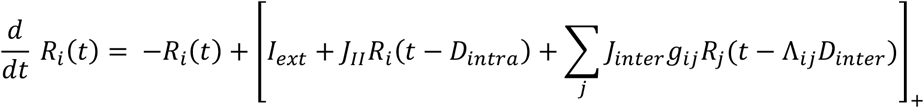

The connectome weights *g* and tract length matrices Λ used for the ING simulations in Figure 4 involve 66 regions within a standard parcellation scheme (Desikan et al., 2006). The matrices are derived from Diffusion Spectrum Imaging (DSI) and averaged over five healthy subjects, as previously published and detailed in Hagmann et al. (2008). The optimal Dynamic Working Point (DWP) providing best correlation with an empirical FC matrix also described in Hagmann et al. (2008) is achieved for *j*_*inter*_ = 6 and *D*_*intra*_ = 1, maintained constant through all simulations with changing *j*_*II*_ and *D*_*inter*_ For the ING large-scale model we phenomenologically emulate a “demyelinating” disease by increasing the delay *D*_*inter*_ of inter-regional coupling. We then restore FC by increasing *j*_*II*_ within each region of the model. The degeneration path of Figure 4D starts at the DWP and ranges from 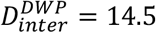 to 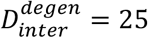. The delay is then maintained at the degeneration value but local inhibition boosted from 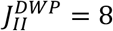 to 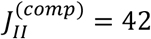 along the compensation path. We generate a simulated FC matrix for 150 points over both paths and project these FCs a bidimensional latent space the vectors spanned by the 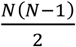 entries of their upper triangular part through a t-Stochastic Neighborood Embedding (tSNE, Hinton & van der Maaten, 2008) projection. A vertical axis is added to Figure 4D to emphasize progression percent along the considered degeneration and compensation paths.

In the large-scale connectome-based PING model, we don’t use delays. However, we introduce multiple inter-regional remote E-to-local E and remote E-to-local I current variables (one per presynaptic neighbor of the considered region. In the dynamic equations of these variables, coupling conductances between regions *i* and *j* are thus further scaled by the relative connectome strength *g*_*ij*_ given by the adopted connectome matrix, i.e. *G*_*EE*_ and *G*_*IE*_ in PING region recurrent currents above are replaced by *G*_*EE*_*g*_*ij*_ and *G*_*IE*_*g*_*ij*_. The architecture of the large-scale model is identical to what described by Benitez Stulz et al. (2023; see previous chapter in this thesis), with all unspecified parameters identical to what indicated therein. The model displays phase transitions as an effect of changing *G*_*EE*_, *G*_*IE*_ and the external drive *I*_*ext*_ to each region. Specifically, the level of overall synchronization in the system varies as a function of parameters of global and local dynamics. Following Benitez Stulz (2023), we select reference DWPs not far away a critical line of transition to global synchronization as tracked by the *Phase-Locking-Value* (PLV; Lachaux et al. (1999)). Specifically, in Figure 5, we choose a slightly subcritical DWP with values 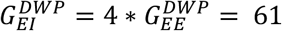 and 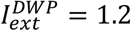. To evaluate synchrony, we first extract the instantaneous phases of oscillation *ϕ*_i_(*t*) of each region *i*. For every pair of regions we evaluate then a pairwise locking index:

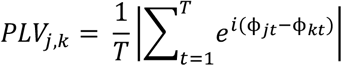

and then compute an overall global PLV by averaging over all possible links:

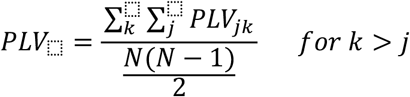

The used connectome matrix, weighed and directed, has been obtained using retrograde tracing experiments in Non-Human Primates (Markov et al., 2015) and include all existing connections between a subset of 29 regions. For the PING large-scale model we phenomenologically emulate “connectivity loss” in a degenerative disease by reducing, in Figure 5, the strength *G*_*IE*_ of inter-regional dysynaptic inhibitory coupling. We then restore FC by increasing *j*_*EI*_ within each region of the model.

The degeneration path of Figure 5C starts at the DWP and ranges from 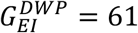 to 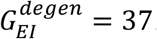. The coupling strength is then maintained at the degeneration value but local I to E coupling increased from 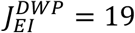 to 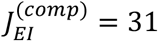 along the compensation path. Once again, we generate a simulated FC matrix for 150 points over both paths. We then study, in Figure 5C, how similar the obtained FC matrices are to the reference FC_DWP_ at the DWP point and to FC_degen_ at the end of the degeneration path, using Pearson Correlation between the upper-triangular part vectors of the matrices as metric of similarity. In Figure S5, we compensate degeneration of long-rang disynaptic inhibition by raising the level of background driving current. The degeneration path ranges from 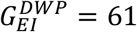 to 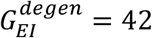. Along the compensation path, the background drive is increased from 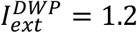 to 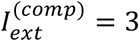. Finally, in Figure S6, we compensate plasticity of long-range excitation by raising the local E-to-I coupling. The plasticity change path ranges from 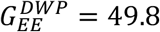 to 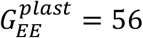. Along the homeostatic compensation path, the local E to I coupling is increased from 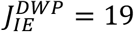 to 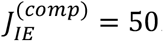.

### Numerical Integration

Simulations of the large-scale ING model are conducted within a custom-made C code, available upon request, using a fourth-order Runge-Kutta scheme of integration, with integration step *τ*_int_ = 0.001 in arbitrary units. Simulations of the large-scale PING model are conducted using the Virtual Brain simulation environment (https://thevirtualbrain.org) to which a DumontGutkin neural mass class (corresponding to our PING ERM) has been added in the latest releases. The Heun integration scheme is used with integration step *τ*_int_ = 0.0001 in arbitrary units.

Note that all simulations (toy- and large-scale) are exactly deterministic (for both the ING and PING cases) and that, therefore, signal irregularity eventually arises through inter-regional interactions leading to quasi-periodicity and chaos.

### Macroscopic Phase Response Curves of PING and ING models

The (infinitesimal) Phase Response Curve (PRC) is defined mathematically for infinitesimally small perturbations to a trajectory unrolling on a limit cycle *x*_*cycle*_(*t*), and can be computed rigorously via the so-called adjoint method (Brown et al. 2004). Specifically for a limit cycle solution evolving according to the equation:

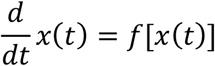

then the PRC obeys the defining equation:

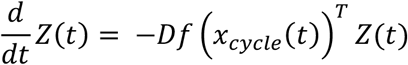

where *Df*(⋅)^*T*^ is the transpose of the time-dependent Jacobian matrix of the system *f*. Note that, since the considered trajectory is a limit cycle, it is periodic so it can be expressed in term of a phase *ϕ* = *2πt* mod *T*, where *T* is the period of the considered oscillation. The PRC can thus be seen as the (phase-dependent) phase advancement or delay induced by applying an infinitesimal perturbation at a given phase of the limit cycle.

For the PING system, the PRCs shown in Figure 2 and Figures S1-S3 are evaluated by direct numeric integration of the adjoint equation of the macroscopic ERM equations. Details are provided in Dumont & Gutkin (2019). We also performed numerical experiments on spiking networks of QIF neurons to which the PING ERM is equivalent. There we applied a short (2 time-steps) square wave of additional external stimulation (+20% in drive) to each excitatory neuron in the spiking network at different phases of the simulated collective oscillation and measured the resulting displacement of the periodic events of synchronized firing in the corresponding raster plot (random seed generators were stored to regenerate the same quenched disorder in both the unperturbed and perturbed QIF network simulations). The obtained stimulation-phase dependent collective population-level phase shifts were then rescaled to match the variation range of the ERM-based PRC, for the sake of a better comparison in Fig. 1D.

For the ING model, the piecewise linear nature of the differential equation describing the macroscopic collective activity allows a direct analytic evaluation of the PRC curve. Details of derivation are too long to be reported here but are described in full detail in the supporting information of Battaglia, Brunel and Hansel (2007). Therein we prove that the PRC of the ING dIReLU model can be written as:

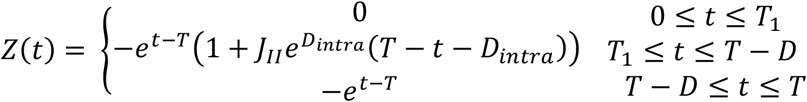

where *T* is the oscillation period and *T*_1_ the time within the period at which the total input current to the unit *I*_*tot*_ = *I*_*ext*_ + *j*_*II*_*R*(*t − D*_*intra*_) crosses the threshold for firing becoming positive from negative. The values of *T* and *T*_1_ can be obtained by numerically solving this system of nonlinear equations:

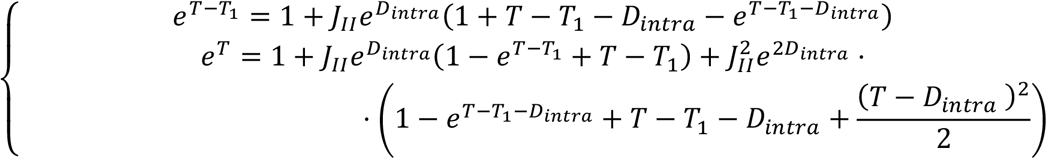

All PRCs shown in Figure 3 are obtained solving and evaluating the above expressions. We used a standard Levenberg-Marquardt solver for finding the roots of the above nonlinear system.

### Determination of stable phase differences between coupled regions

The evolution of the phase-difference Δϕ(t) = *ϕ*_1_(*t*) – *ϕ*_*2*_(*t*) between two reciprocally coupled oscillators “1” and “2” can be evaluated in terms of the PRC curve, according to the weakly coupled phase oscillator theory (Kuramoto, 1991; Pikovski et al., 2003):

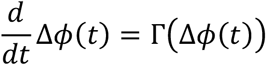

Here Γ(Δ*ϕ*) can be expressed as the difference between two convolution integrals, describing respectively the integrated action of inputs *s*_1_(*t*) (or *s*_*2*_(*t*)) received by “1” (or “2”) from “2” (or “1”) on the phase of “1” (or “2”). The phase *ϕ*_1_ is shifted to a value *ϕ*_1_ + Γ_1_(Δ*ϕ*), where:

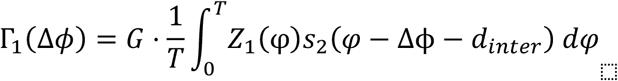

Indeed, the limit cycle waveform of the input *s*_2_ can be seen as the continuous sum of instantaneous pulses, each shifting the phase of *ϕ*_1_ of an infinitesimal amount proportional to the PRC of “1” at the phase *φ* at which perturbation is received and to the perceived stimulus intensity, which is the value of the input *s*_*2*_ at the phase *φ*, shifted of the current phase shift Δ*ϕ* and of an additional shift 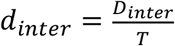 mod *T* linked to the delay of interaction. Here *G* is the strength of the considered inter-regional coupling. Analogously, the phase *ϕ*_*2*_ is shifted to *ϕ*_*2*_ + Γ_*2*_(Δ*ϕ*), with:

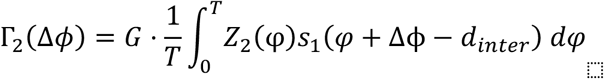

so that the overall rate of variation of Δ*ϕ* is given, by the difference between the phase shifts of “1” and “2”, i.e. :

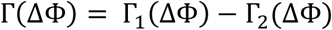

The zeroes of Γ(Δ*ϕ*) thus provide the possible equilibrium values of Δ*ϕ*, stable (or unstable) depending on the negative (or positive) sign of the slope of Γ(Δ*ϕ*) at these zero crossing points. After evaluating the Γ function, the position of its zeroes can again be determined using a standard Levenberg-Marquardt nonlinear solver, thus determining the possible phase-locking differences between “1” and “2”.

Note that, for the ING dIReLU model, the waveforms *s*(*t*) can be exactly derived in analytical form, as reported by Battaglia, Brunel and Hansel (2007):

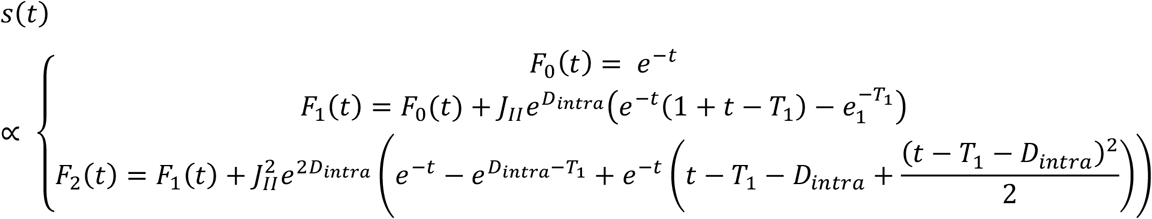

where the three expressions *F*_0,1,*2*_(*t*) hold for the three time-intervals, respectively: 0 ≤ *t* ≤ *T*_1_; *T*_1_ ≤ *t* ≤ *T − D*_*intra*_; and *T − D*_*intra*_ ≤ *t* ≤ *T*, and where, as above, *T* corresponds to the oscillation period and *T*_1_ to the intra-period time at which total inputs cross the threshold of the rectified linear transfer function (*I*_tot_ = 0).

For the PING ERM model, the waveforms to be plugged in the definition of the convolution integrals Γ_1_ and Γ_*2*_ are obtained through numeric integration of the corresponding differential equations.

## ACKNOWLEDGMENTS

S.B., G.D., B. G and D.B were supported by the French Agency for National Research “ERMUNDY” (ANR-18-CE37-0014). S.B., S.C. and D.B. were supported by the French Agency for National Research “HIPPOCOMP” (ANR-18-CE37-0014). S.C and D.B were also supported by the University of Strasbourg Institute for Advanced Studies (USIAS-2020-044), where part of this research has been conducted. We would like to thank Viktor Jirsa and Spase Petkoski for discussions about dynamic compensation and Giovanni Rabuffo for precious help in simulation implementations.

## AUTHOR CONTRIBUTIONS

S.B. and G.D. performed analyses and simulations of toy models, S.C. performed analyses and simulations of large-scale models, B.G., G.D. and D.B. conceived the study, all authors interpreted the results and wrote the manuscript.

## SUPPORTING FIGURES

**Figure S1.**
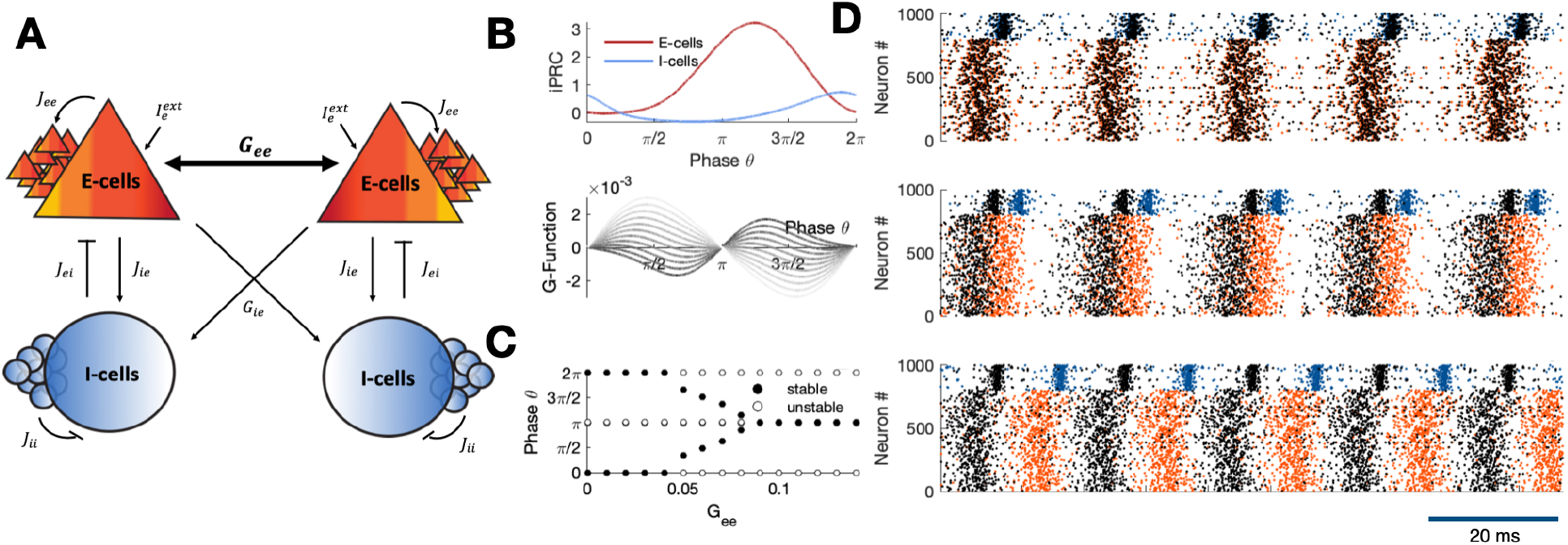
Effects of changing inter-regional E-to-E coupling among PING regions. **(A)** As an effect of modifying *G*_*EE*_, **(B)** the Γ function ruling the dynamics of instantaneous inter-regional phase differences in the ERM model (bottom) is affected but not the local PRC of each population within the region. **C)** By increasing *G*_*EE*_, the initial in-phase locking configuration is destabilized toward out-of-phase and, then, anti-phase locked configurations. **D)** A similar phenomenology is captured by simulating spiking networks of QIF neurons and reducing the strength of E-to-E long range synapses.

**Figure S2.**
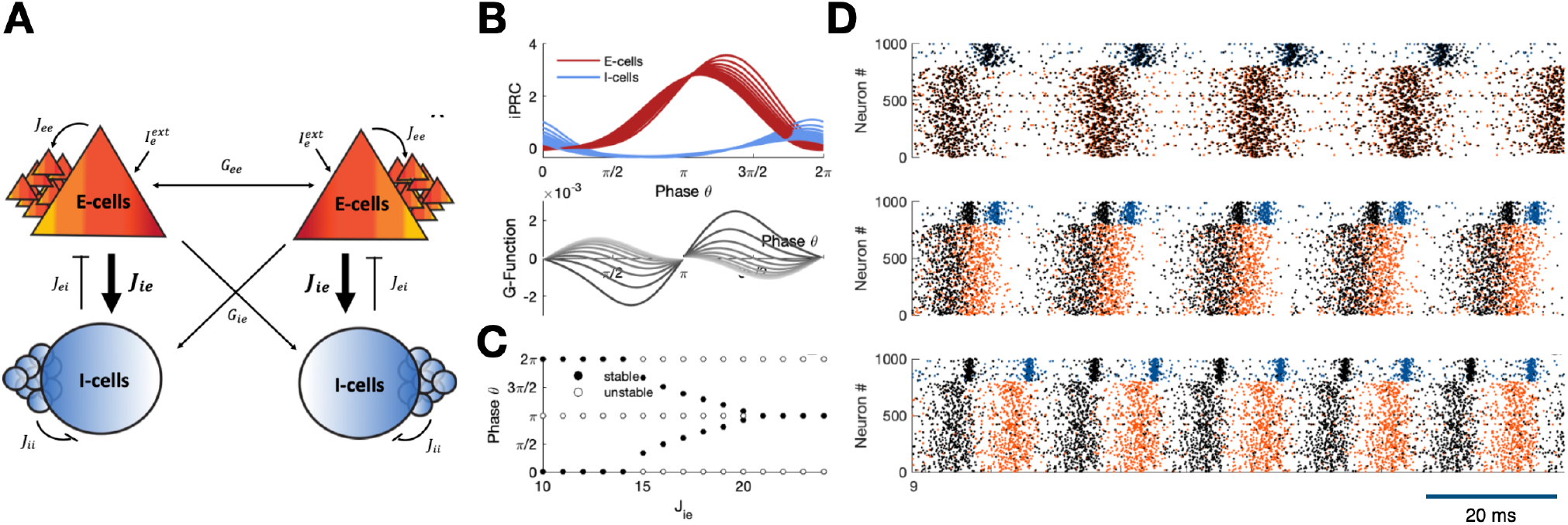
Effects of changing within region E-to-I coupling among PING regions. **(A)** As an effect of modifying *J*_*IE*_, **(B)** the Γ function ruling the dynamics of instantaneous inter-regional phase differences in the ERM model (bottom) is affected, as well as the local PRC of each population within the region. **C)** By increasing *J*_*IE*_, the initial in-phase locking configuration is destabilized toward out-of-phase and, then, anti-phase locked configurations. **D)** A similar phenomenology is captured by simulating spiking networks of QIF neurons and reducing the strength of E-to-I local recurrent synapses.

**Figure S3.**
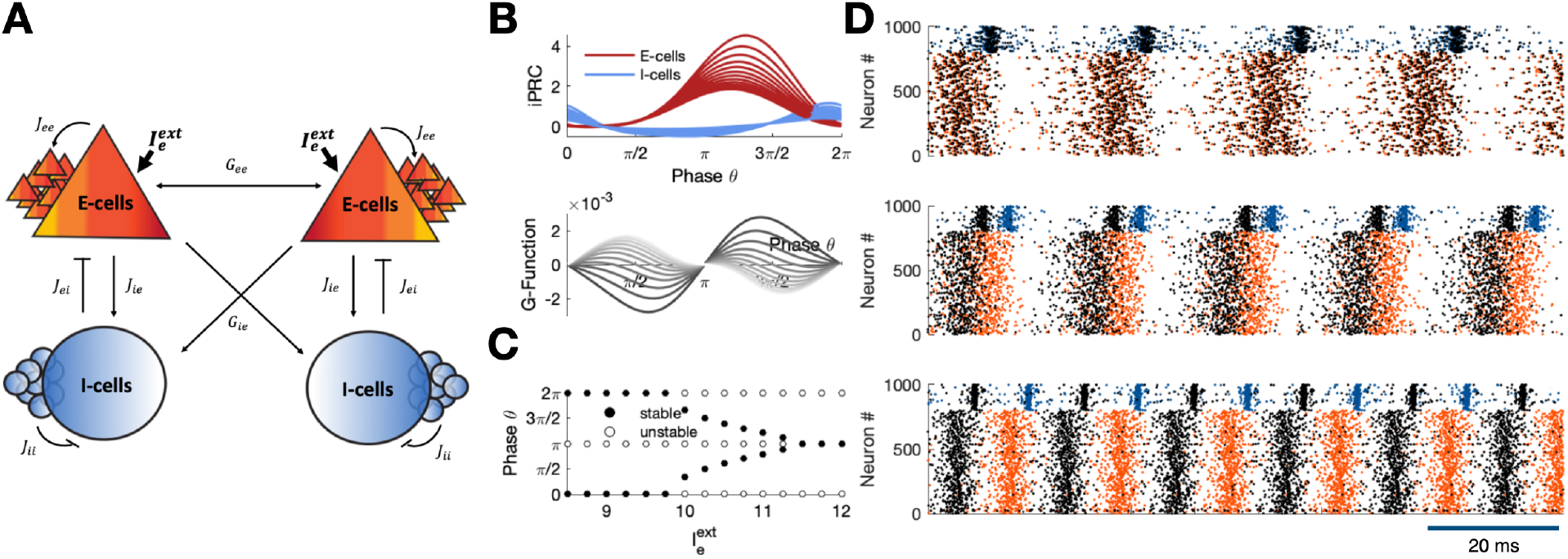
Effects of changing drive to E populations within coupled PING regions. **(A)** As an effect of modifying 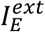, **(B)** the Γ function ruling the dynamics of instantaneous inter-regional phase differences in the ERM model (bottom) is affected, as well as the local PRC of each population within the region. **C)** By increasing 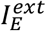, the initial in-phase locking configuration is destabilized toward out-of-phase and, then, anti-phase locked configurations. **D)** A similar phenomenology is captured by simulating spiking networks of QIF neurons and increasing the input current to E neurons.

**Figure S4.**
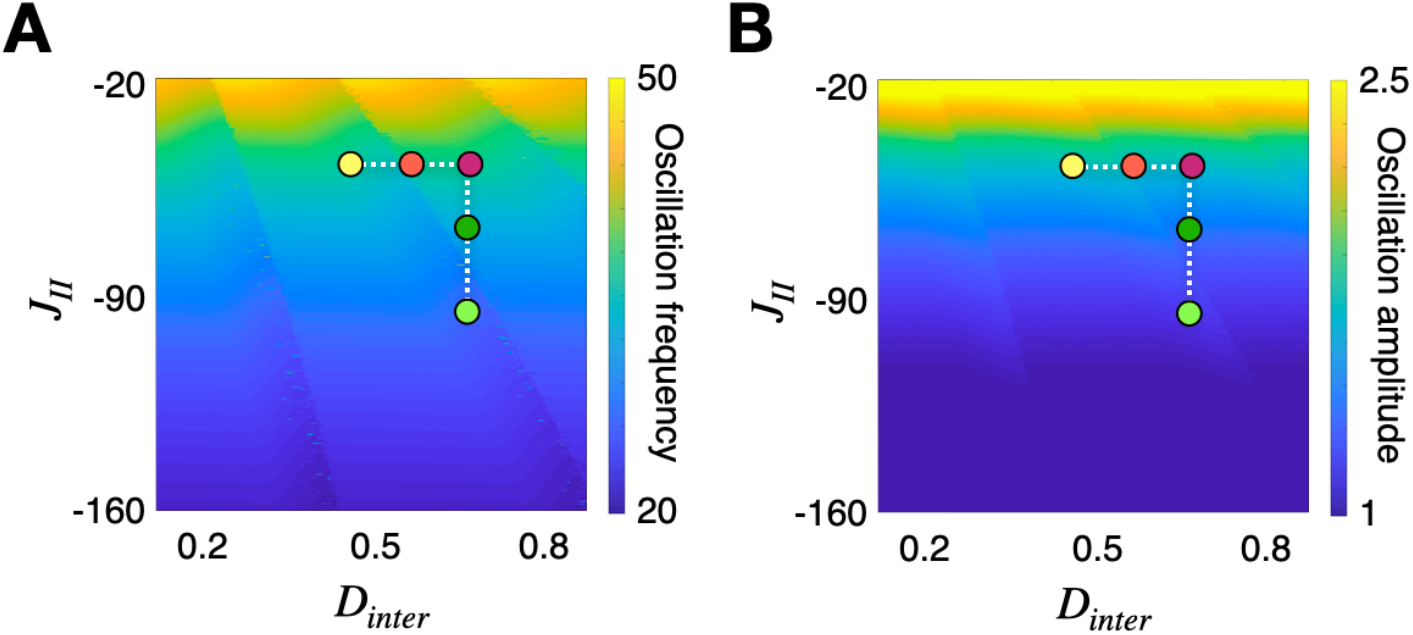
Parameter dependency of the oscillation frequency and amplitude in coupled ING regions. Systematic analysis of the collective oscillation **(A)** frequencies and **(B)** amplitudes obtained for changing *D*_*inter*_ and *j*_*II*_. Highlighted on the surfaces are the same six example configurations as in Figure 3G.

**Figure S5.**
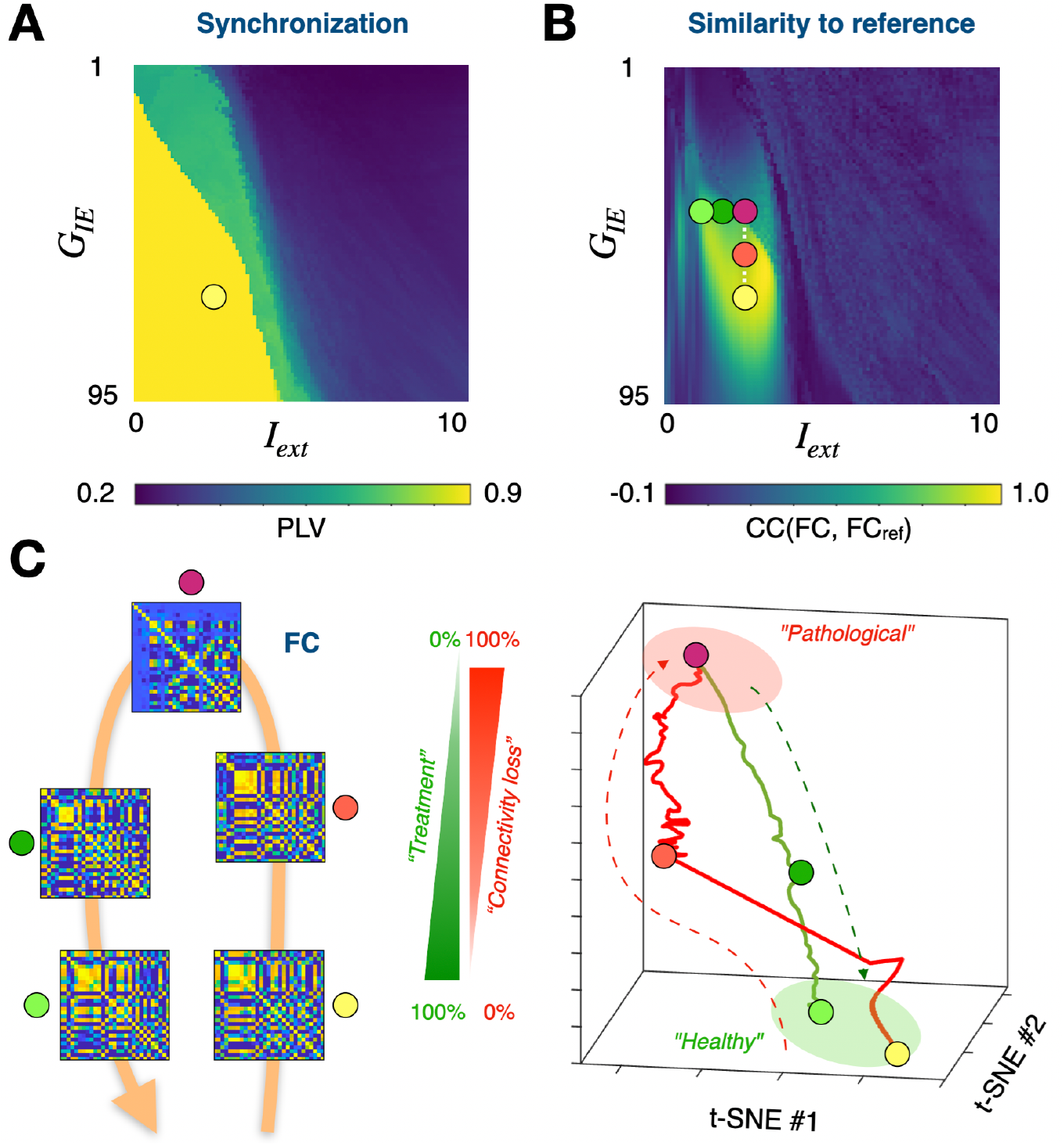
Additional degeneration compensation strategies for a whole-brain model with PING dynamics: adjusting regional drive level. Using the same PING connectome-based model of Figure 5 we consider alternative ways to compensate for structural connectome degeneration by modifying local dynamics. **A)** Dependency of model synchronization level from the strength of inter-regional coupling (disynaptic inhibition) and the level of background regional drive. We consider as reference a Dynamic Working Point (in yellow) close to a transition between synchronous and asynchronous dynamics (but this time on the supercritical side, unlike in Figure 5, showing robustness of the compensation concept, holding for alternative DWPs). (**B**) Similarity between the obtained FC and the reference FC_ref_ matrix as a function of changing DWP. Simulated connectivity loss (decrease of *G*_*IE*_) causes the system to leave this regime along a degeneration pathway. However, increasing *I*_*ext*_ within each region (as in Figure S3) causes the system to follow a compensation pathway entering again a zone of higher global PLV. **(C)** To the left are shown FC matrices obtained at the points along degradation and compensation pathways highlighted in panel **B**. Also shown to the right is a dimensionally reduced representation of FC matrix changes along these pathways, obtained via a non-linear distance preserving t-SNE projection of the sequences of simulated matrices (the vertical dimension reports % progression along the degeneration and compensation pathways).

**Figure S6.**
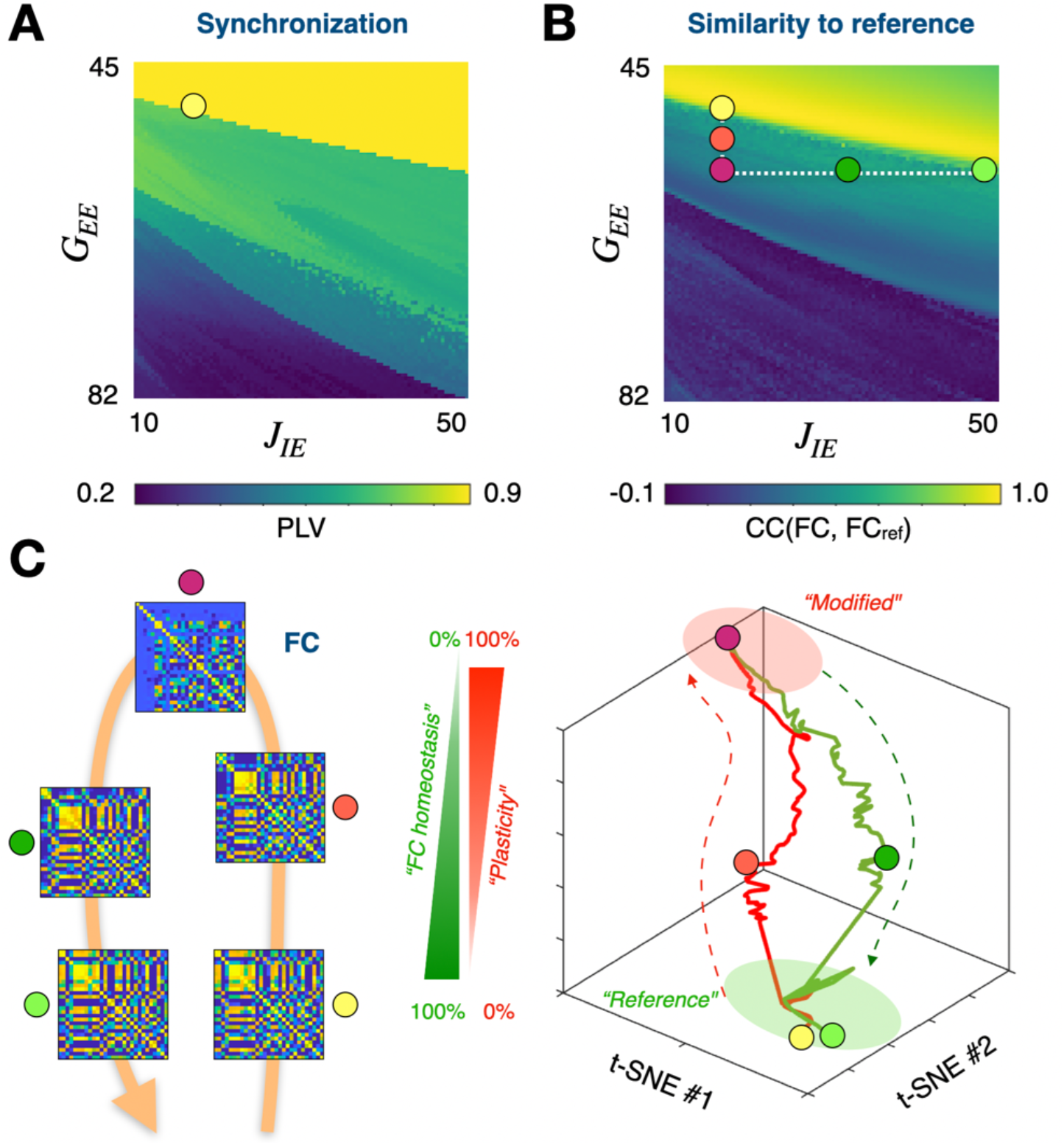
Additional compensation strategies for a whole-brain model with PING dynamics: adjusting local excitation to inhibition strength. Using the same PING connectome-based model of Figure 5 we consider alternative ways to compensate for global dynamics effects of modifying structural connectome by modifying local dynamics. **A)** Dependency of model synchronization level from the strength of (excitatory) inter-regional coupling and the strength of local excitation to inhibitory population. We consider as reference a Dynamic Working Point (in yellow) close to a transition between synchronous and asynchronous dynamics (slightly supercritical). (**B**) Similarity between the obtained FC and the reference FC_ref_ matrix as a function of changing DWP. Simulated connectivity plasticity (increase of ⬚_*EE*_) causes the system to leave this regime modifying inter-regional phase relations (as in Figure S1). However, increasing *J*_*IE*_ within each region (as in Figure S2) causes the system to implement FC homeostasis entering again a zone of higher global PLV. **(C)** To the left are shown FC matrices obtained at the points along degradation and compensation pathways highlighted in panel **B**. Also shown to the right is a dimensionally reduced representation of FC matrix changes along these pathways, obtained via a non-linear distance preserving t-SNE projection of the sequences of simulated matrices (the vertical dimension reports % progression along the plasticity and homeostasis pathways).

## Notes

**Conflicts of interest** The authors declare no competing interest.

### Competing Interest Statement

The authors have declared no competing interest.

